# Emergence of tolerance and avoidance strategies from local and systemic responses to nitrogen: insights from modelling of auxin-mediated root plasticity

**DOI:** 10.64898/2026.09.08.750047

**Authors:** Jie Lu, Jin Wang, Alejandro Morales, Jochem B Evers

## Abstract

- Understanding how root phenotypic plasticity enhances resource-use efficiency can help understand the outcomes of plant competition and identify suitable genotypes for medium- to low-input agricultural systems. Auxin regulates multiple root growth processes, including the root architectural responses to nitrogen (N). We explored the extent to which auxin-mediated local and systemic responses to external and internal N influences plant N uptake and use, using a functional-structural-plant (FSP) modelling approach.
- A simplified auxin module was developed at the level of the organ and integrated into an FSP model to represent physiological plastic root responses to N. Model performance was evaluated against experimental data. We then ran simulations under various N conditions with local or systemic responses enabled or disabled, to quantify their contribution to N uptake and use.
- Simulations showed that local auxin responses enhanced N uptake by distributing more roots towards deeper soil layers and increased N forage, thereby avoiding N stress. Systemic auxin responses reduced N uptake by distributing more roots near the top soil layers, which reduced plant size and N demand, thereby enhancing stress tolerance. This points to a trade-off in N uptake between the tolerance and avoidance strategies, which can be traced back to biomass and N investment resulting from source-sink relationships in the plant.
- Representing hormone-mediated root plasticity in an FSP model provides mechanistic insights into plant strategies for resource capture.

## Introduction

In order to lower crop N inputs to mitigate environmental pollution, crops that take up and use nitrogen (N) efficiently are essential. This efficiency can be improved by plastic responses in plant development to low N availability. Plastic responses, both locally (organ-level) and systemically (whole-plant level), to internal or environmental signals, allow the plant to deal with the deficiency (Alvarez et al., 2012; De Kroon et al., 2009). Low soil N availability in the direct vicinity of a root can trigger fast growth responses of the root. In addition, internal signals such as plant N status, which is the result of the uptake of soil N, can elicit systemic plant responses, such as a change in biomass allocation, resulting in a higher root/shoot biomass ratio (De Kroon et al., 2009). Many of these local and systemic responses to internal and environmental signals are regulated through plant hormone-dependent mechanisms (Eckardt, 2015; Zandalinas et al., 2020). The eventual root system architecture and the associated uptake of N during plant development are thus functions of local and systemic responses to N in the soil and in the plant. How systemic and local responses to N contribute to plant survival strategies remains unclear.

Plants differ in the strategies they use to minimize the negative effects of environmental stress (Grime, 1974). Stress avoidance and stress tolerance are two of the major plant survival strategies. In general, avoidance reduces the impact of environmental stress, while tolerance enables plants to withstand and maintain function under stressed conditions (Puijalon et al., 2011). For instance, the plastic response of increased root elongation rate, allowing root foraging for soil resources to reduce the impact of environmental stress, is a typical avoidance strategy (Wang et al., 2020). An increase in specific leaf area to optimize light capture to survive under shaded conditions is a typical tolerance strategy (Gommers et al., 2013). Stress-tolerant plants were usually associated with a slower growth rate than other species (Kitajima, 1994). Clearly distinguishing traits relevant to tolerance and avoidance strategies is useful for advancing the mechanistic understanding of plastic responses to stress for crop improvement.

Auxin is an important plant hormone influencing a multitude of processes related to plant growth, development, and responses to various stresses (Vanneste et al., 2025). Auxin is produced in both root and shoot meristems. Both shoot- and root-produced auxin controls developmental processes in the roots. Shoot-produced auxin is transported to the root meristems and controls lateral root formation, root elongation, and root orientation (Du & Scheres, 2018; Vanneste et al., 2025). Root-produced auxin mainly participates in lateral root initiation, root elongation, and root orientation in root tips (Du & Scheres, 2018; Liu & von Wirén, 2022; Meier et al., 2020; Vanneste et al., 2025).

The presence of N in the soil or within the plant affects auxin distribution within each single root, and therefore affects root system architectural development (Jia et al., 2022). Low external N in the form of nitrate enhances auxin biosynthesis and auxin efflux in both axial and lateral root meristems (Jia et al., 2023; Jia & von Wirén, 2020; Krouk et al., 2011). More auxin produced in the shoot can be transported to the root when plant N status is low (Asim et al., 2020; Giehl & von Wirén, 2014; Tian et al., 2008). In this study, we defined the auxin transported from the shoot to the root tips, regulated by the plant’s internal N status, as the systemic response. Meanwhile, the part of auxin produced by root tips and also the efflux of auxin in the root tip regulated by external soil N concentration were defined as local responses.

In this study, we address the question of to what extent root system architecture is determined by systemic and local root responses to N availability mediated by auxin, and how this affects plant N use in relation to plant stress avoidance or tolerance strategies. Disentangling the relative contributions of systemic and local responses to root system architecture experimentally is challenging, even when using mutants. For instance, the pleiotropic effect of a gene may include lethal outcomes, masking the changes in specific phenotypes (Tonsor et al., 2005).

On the other hand, in mathematical modelling mechanisms and processes can be toggled and tested at will, making it a useful approach to help disentangle complex mechanisms interacting from the plant level to cellular levels, and evaluating their effects on plant function and development (Grieneisen et al., 2007; Lu et al., 2024b; Postma et al., 2014; Postma & Lynch, 2011; van den Berg et al., 2021). Mathematical modelling of cell-level auxin regulation in the root meristem has been developed to address multiple fundamental questions, such as the effect of auxin reflux on lateral root priming or the role of PIN proteins on polar auxin efflux (Grieneisen et al., 2007; Rutten et al., 2022; van den Berg et al., 2021). Due to the complexity in the interplay between auxin activity, plant functioning, and environmental signals, cell-level auxin modelling that is limited to the root meristem cannot efficiently address questions related to whole root-system architecture formation and soil resource capture. In contrast to cell-level modelling, functional-structural plant (FSP) models simulate feedback between physiological processes and plant architecture and the effects on resource capture at the organ and plant level, rather than the cell level, and plant growth is simulated as a function of environmental factors (Boer et al., 2020; Vos et al., 2010). FSP models can be used to represent physiological processes at the level of the plant organ, as has been done for auxin mechanisms in the shoot (Prusinkiewicz et al., 2009), but FSP models have not yet been used to address auxin-driven root responses to soil and plant N status and its consequences for root architecture and N uptake. The objective is to identify the role of local and systemic responses in how the root system architecture responds to soil N and the consequences for plant growth, using a simulation model that simulates root architectural plasticity based on auxin physiology in response to nitrogen.

## Methods

The goal of our modelling approach was to capture plastic root responses to nitrogen using a simple physiological basis that goes beyond mere signal-response curves, in order to simulate root system architecture as a function of N. Therefore, rather than adopting and developing an elaborate molecular-physiological model of root elongation or side root formation at the cell-level (Dyson et al., 2014; Péret et al., 2013), we chose to focus on a crucial determinant of root architectural development, auxin (Vanneste et al., 2025). We considered auxin physiology at the level of the root organ, similar to the level of detail of the approach followed in Prusinkiewicz et al (2009). We first developed a module (a model component) that links auxin production and allocation of shoot- and root-derived auxin to the root meristems, affecting lateral root formation and root elongation as functions of internal and external N levels; we do not consider root angle as a function of auxin activity. Then, we integrated this module into an FSP model of wheat root and shoot growth, using local soil N and plant N as inputs to the module. Subsequently, to evaluate model performance on plant growth, root profile, and N uptake, we ran simulations under several N scenarios and compared the output with observations from an experiment with different N levels on wheat. Finally, we addressed our research question by conducting virtual experiments exploring the effects of both the systemic and local root responses to internal and external N on root system architecture, N uptake, biomass, and plant N concentration.

### Model description

#### FSP model

The wheat FSP model has been developed in the Virtual Plant Lab (VPL) (Morales et al., 2025) implemented in the Julia programming language. The wheat model simulates plant growth based on carbon and nitrogen sink-source relationships driven by light, temperature and soil N, and it is based on earlier work (de Vries et al., 2021.; Evers et al., 2010; Evers et al., 2007; Lu, et al., 2024a) but modified to account for responses to N availability.

The model increases biomass allocation to the roots under low N based on the functional-equilibrium theory (Shipley and Meziane, 2002). This is achieved with a plasticity factor Λ_root:shoot_ (Lu et al., 2024b; eq. S4) that multiplies the root sink strength in the model in response to low plant N. Tillers were included to represent the characteristic shoot architecture of wheat. Tiller production is determined by the carbon sink-source ratio (Evers et al., 2010). To account for the variation of tiller number caused by environmental factors such as shading (Shang et al., 2021), we introduce a fixed probability to determine the final tiller number of a wheat plant (Wang et al., 2025). We assumed that the effect of auxin-mediated root architectural plasticity influences total shoot biomass per plant and not the distribution of biomass across tillers. In addition, N uptake becomes insensitive to the number of axials roots for values above 10 (Lu et al., 2024b, Lu et al., 2025). For these reasons, tiller number was not linked to axial root production in the current model. The architectural development of the wheat roots is based on ArchiSimple with monocot characteristics (Pagès et al., 2014a, 2014b). To simulate N-uptake, high and low-affinity transport kinetics are used (Lu et al., 2024b). Soil is represented by a 3D grid with voxels of 10×10×10 cm, with N distribution within each voxel assumed to be uniform. Also, to help address the research question without interference from soil N dynamics other than N uptake, N redistribution in the soil was not simulated. A more detailed description of the model and relevant equations is available in the Supplement (“FSP model description” section).

#### Auxin module

The auxin module was constructed based on current experimental evidence on auxin biosynthesis, transport, and regulation in roots to investigate their combined effects on root architectural development at the whole plant level. For physiological parameters, whenever possible, values were used from literature on wheat. Where quantitative information was unavailable, biologically plausible assumptions were made based on published observations, since the functioning of auxin is conserved across dicot and monocot species such as Arabidopsis or maize (Kiba et al., 2011; Li et al., 2014).

We implemented three main modes of action for auxin in our model: 1) auxin drives the formation of lateral root primordium in pericycle tissues; 2) shoot-derived auxin directly influences root emergence and 3) elevated auxin levels in cell division and elongation zones suppress root elongation. The module and variable definitions are summarized in Fig. 1 and Table S1.

**Figure 1.**
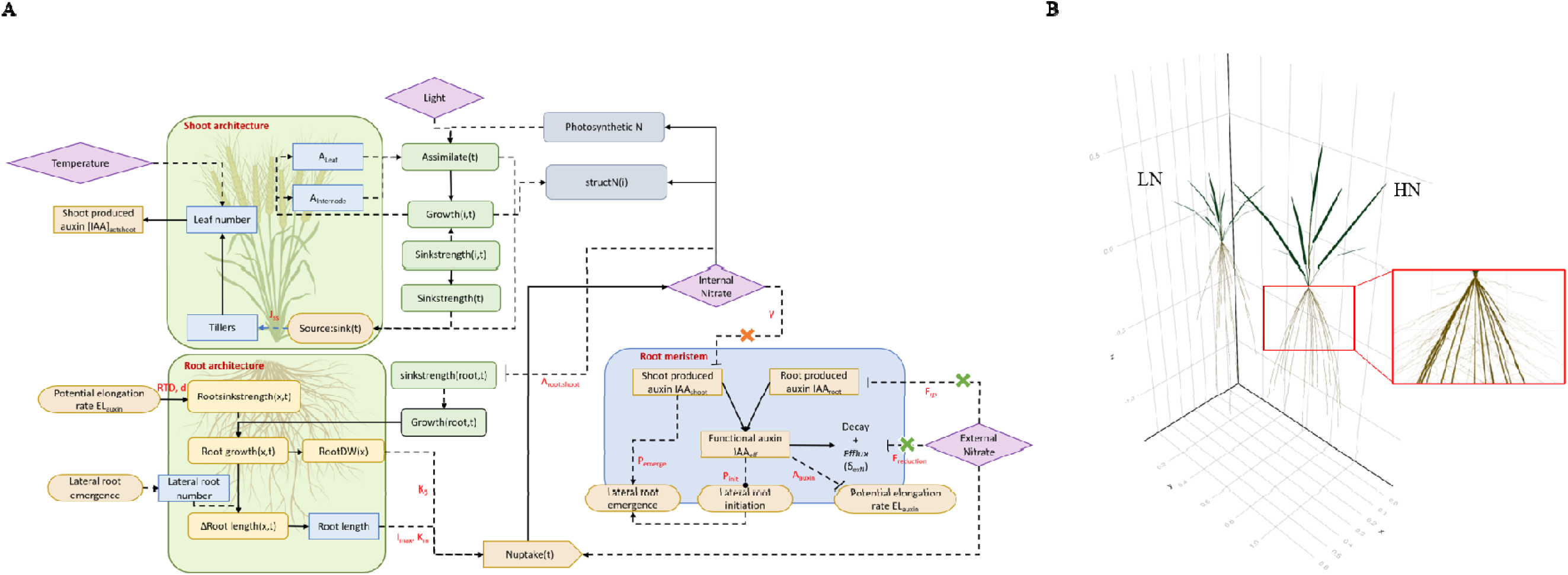
Model diagram of auxin functioning in the root apical meristem in response to nitrogen (Nitrate, Panel A) and model graphic illustrations under high N (HN, 3000 µmol/L) and low N (LN, 10 µmol/L, Panel B). In Panel A, the solid lines represent matter flows and the dashed lines represent the information flows. The diamonds represent environmental factors. The rectangles with round edges represent C (green) and N (grey) sinks or sources at organ level (leaf, internode, grain and root system). The yellow rectangles with round edges represent C sink or source at individual root level. The blue rectangles represent plant morphological traits. The light brown rectangles represent different source of auxin. The ellipses represent intermediate model variables. A regular arrowhead represents a positive relation; a flat arrowhead is a negative relation, and a dot arrowhead represents a positive effect at low values but a negative effect at high values. Subscripts: *t* represents the time unit; *i* represents plant organ (i.e. root system, leaf, internodes, grain); and x represents root segment. The red symbols represent model parameters relevant to the relationship they are linked to in the figure. The orange cross represents the disabled relationship in simulation scenarios in which only local responses were enabled. The green crosses represent the disabled relationship in simulation scenarios in which only local responses were enabled. The green crosses represent the disabled relationship in simulation scenarios in which only systemic responses were enabled. Panel B: illustrations of plant architecture under low or high N from simulations at day 40, with the insert zooming in to show the lateral roots.

Two main auxin sources are distinguished and independent (shoot-produced and root-produced auxin), and these are involved in different aspects of root development (Casimiro et al., 2001; Reed et al., 1998). For simplicity, the model assumes axial root meristems and shoot meristems were the same size. Auxin levels in meristems are expressed in arbitrary units as a concentration. In this study, lateral root initiation and emergence, and potential root elongation rate are directly linked to auxin level (Fig. 1). To scale auxin functions from the root-tip level to the root-system level within a whole-plant FSP model, the root cap and the zones of cell division, elongation and maturation were not represented as separate spatial compartments. Instead, their functions were integrated into a root-tip functional unit, represented in the model by the root apical meristem. Auxin regulates the potential elongation of the root meristem, while actual root elongation is determined by C source–sink relations and represented by the production of fixed-length root segments. All parameter values are listed in Table S2 and Table S3.

### Auxin production and decay

Auxin biosynthesis happens in shoot apices and leaf primordia (Ha et al., 2010). Meanwhile, shoot-produced auxin mainly contributes to root formation at the early growth stage (Bhalerao et al., 2002). Therefore, the potential contribution of shoot auxin production to auxin concentration in the root meristem (IAA_s,pot_, expressed in arbitrary auxin units (a.u.) per day) is assumed to be a constant per leaf for the four youngest leaves of a tiller, as long as a leaf is younger than 8 days (Bhalerao et al., 2002)).

There is an interaction between auxin transport and assimilate transport to the root. Sugar transport to the root (such as sucrose) can influence auxin long-distance transportation by enhancing the expression of auxin-transportation-related genes, such as PIN1 (Lin et al., 2016; Sakr et al., 2018; Zhang et al., 2020). In addition, Balasubramanian et al. (2024) linked modulation of polar auxin transport with carbon source–sink relationships and carbon sink strength. Both studies indirectly suggested that organs with stronger carbon demand attract more assimilates and auxin. Therefore, in the model, we assume that when the shoot-produced auxin is transported to the root, it redistributes to each root meristem based on the relative carbon sink strengths (*rss_root_*, eq. 1) of each root every model time step of one day.

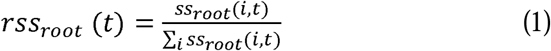

The *ss_root_* represents root sink strength (eq. S3) at a given daily step (*t*) for each root (*i*).Next to the shoot, auxin synthesis also happens in root apices (Roychoudhry et al., 2021). Our model assumes a constant potential auxin production in the axial and lateral root meristems under high N conditions (*IAA_r,pot_*, a.u. per day). The auxin concentration in the axial and lateral root meristems has been reported to scale approximately with the ratio of axial to lateral root diameter (Liu et al., 2010). Based on this observation, in the model, we assumed that the ratio of *IAA_r,pot_* between axial and lateral roots is equal to the ratio of their diameters. Since the sizes of the shoot and root meristem were assumed to be identical, the shoot-produced auxin concentration adds to root-produced auxin concentration in each root meristem (eq. 2) to calculate the increase of total auxin concentration (*IAA_total_*, a.u.) in each root meristem, which potentially has an influence on root elongation and lateral root initiation.

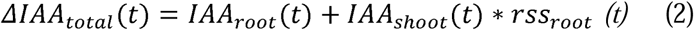

*IAA_root_*represents the rate of auxin production in the root at given external N conditions in each root meristem and *IAA_shoot_* represents the rate of auxin production in the shoot at given plant N concentration in each root meristem.

In the model, we combined the rate of auxin decay in the apical root meristem and the rate of auxin transport out of the meristem in response to external N (δ*_exN_*, per day, eq. 13), as in both cases, auxin has no further influence on lateral root emergence or elongation. This results in the ‘effective auxin’ (*IAA_eff_*, a.u. per day), the auxin that stays in each root meristem after accounting for auxin efflux:

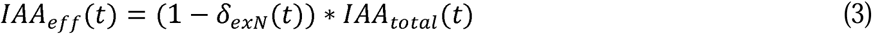

### The function of shoot-produced auxin

Lateral roots emerge from the pericycle of the axial root and start to produce auxin after the lateral root meristem is formed (Péret et al. 2009). Shoot-produced auxin is transported through the vascular cylinder to root tips and mainly contributes to lateral root emergence (Bhalerao et al., 2002). In Arabidopsis, when the flux of shoot-derived auxin to the roots is lowered by removing the shoot apical meristem, the number of lateral roots is heavily reduced (Reed et al., 1998). Conversely, when applying extra auxin at the root and shoot junction, the number of lateral roots increases (Fig. S1A). When auxin is added close to the root tip, no significant effect on lateral root density was found in the same study, indicating little effect of auxin on lateral root initiation (Reed et al., 1998). Although experimental studies generally consider shoot- and root-derived auxin together in regulating lateral root development, the present model represents their roles separately by developmental stage. Based on experimental evidence on the site of lateral root emergence and observations from early developmental stages (Péret et al. 2009; Bhalerao et al., 2002), shoot-derived auxin transported through the axial root was assumed to modulate the probability distribution of lateral root emergence. Following the establishment of a lateral root meristem, auxin produced locally within the lateral root was assumed to regulate its subsequent potential elongation rate primarily. The probability that a lateral root will emerge is calculated as a function of the rate of shoot-produced auxin *(IAA_shoot_,* eq. 4, Fig. S1B), for each lateral root primordium initiated (initiation itself is calculated separately; see ‘The function of auxin in root meristem’ below). Since new lateral roots rarely emerge on old axial root segments (Dubrovsky et al., 2012), root emergence occurs when the lateral root primordium appears, based on the probability (*P_emerge_*) of a lateral root emergence.

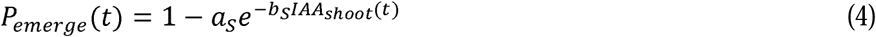

The parameters *a_S_* and *b_S_* are empirical coefficients derived from the literature (Fig. S1; Reed et al., 1998). To align the relative shoot-derived auxin level in the model with the 0–10 μM exogenous auxin range used to parameterize the experimental response curve, simulated auxin values were multiplied by a scaling factor of 10 before being applied to the response function.

### The function of auxin in the root meristem

The shootward transport of auxin through PIN2 transporters in the cortex plays a role in axial root elongation (Liu et al., 2022). Auxin concentration was considered to influence axial root elongation negatively (Reed et al., 1998). In our model, we consider total root auxin in each axial root apical meristem to directly affect the potential root elongation rate (Fig. S2A).

Based on the experiment by Reed et al. (1998), a modifier (*A_auxin_*, eq. 5, Fig. S2B) of the potential root elongation rate was derived as a function of total auxin in the root meristem (*IAA_eff_*) by normalizing with the minimum value equal to 1, which means no effect of auxin on elongation rate. In the model, this modifier *A_auxin_* multiplies the potential root elongation rate parameter (EL, m/m per day, eq. 6).

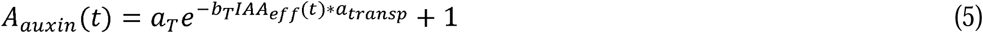

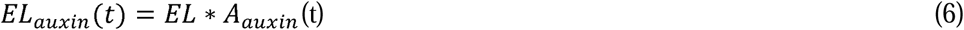

The parameters *a_T_* and *b_T_* are coefficients fitted to data from the literature (Reed et al., 1998; Fig. S2B). As evidence is insufficient to support separate auxin response functions for axial and lateral roots, a common response function was used. A transport coefficient (*a_transp_*=100) was introduced to scale the effective auxin signal in lateral roots to the common response function, in order for the simulated *IAA_eff_* in lateral root tips to exert a meaningful effect on root elongation under the same response parameters (Eq. 5).

When extremely low IAA is transported to cell division and elongation zones, cell activity cannot be maintained (Wang et al., 2023a) and therefore the potential root elongation rate can be reduced. To account for this, a threshold (*V_auxin_*, a.u.) of *IAA_eff_* is included, and below this threshold, we assume the potential root elongation rate is 0.

Auxin in the root meristem promotes lateral root initiation (Dubrovsky et al., 2008; Ivanchenko et al., 2010). However, in Arabidopsis, when auxin application exceeds 12.5 nM, auxin action reverses and starts to suppress lateral root initiation (Ivanchenko et al., 2010).

Based on data from Ivanchenko et al., (2010, Fig. S3), a modified Lambert W function (eq. 7) was derived to represent the probability of lateral root initiation (P_init_) as a function of *IAA_eff_* in the model.

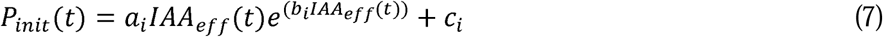

The a_i_ and b_i_ co-determine the steepness and the peak of the curve. The c_i_ determines the probability of the lateral root initiation when auxin is high.

### Auxin translocation in response to internal plant nitrogen

A decreased plant N concentration tends to increase the translocation of shoot-produced auxin to the root system (Asim et al., 2020; Giehl et al., 2014). The auxin concentration in the maize primary root tip is negatively associated with total plant nitrogen concentration (Tian et al., 2008). In the model, we represent the result of auxin translocation as a function of internal plant N. Therefore, a factor to represent the amount of shoot auxin translocated to root (γ, the fraction of auxin at the root transported from the shoot) in response to plant internal N concentration was derived based on the available data points in Tian et al., (2008) and normalized with a maximum value of equal to 1 when total N concentration is equal to *Nc_min_* (eq. 8, 9, Fig. S4B). When the total N concentration is higher than *Nc_max_*, we assume no further decrease in shoot-derived auxin since no further measured information beyond that N concentration was available.

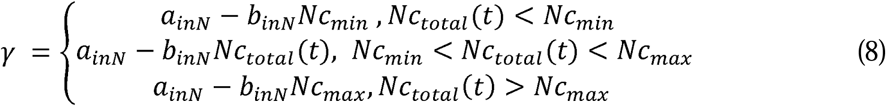

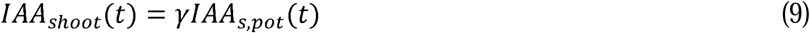

The a_inN_, b_inN_ represents the coefficient, and the value is derived based on Tian et al., (2008).

### Auxin biosynthesis and translocation in response to external nitrate

Low nitrate in the soil around the root tip facilitates auxin biosynthesis in the lateral root tips (Jia et al., 2020). In addition, low external nitrate can promote auxin efflux, such as through NRT1.1 in the lateral root or PIN1a in the main roots (Krouk et al., 2011; Song et al., 2013; Wang et al., 2023b). The decay of IAA in dark conditions is relatively low (Yamakawa et al., 1979), therefore, the decrease in root meristem under low external nitrate conditions is mainly due to auxin efflux. In the model, negative linear relationships are assumed between IAA concentration produced by the root (Fig. S5A) or shootward transportation of IAA from the root tip (Fig. S5B) and external N are derived and the same logic as before is used to normalize the values as the multipliers (Liu et al., 2010, Eq. 10, eq. 11, eq. 12, eq. 13, Fig. S5C, D).

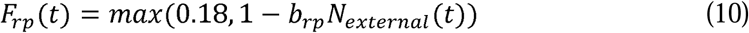

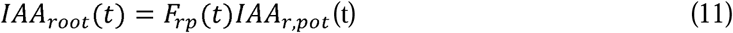

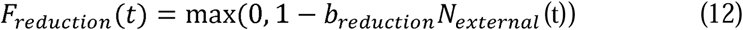

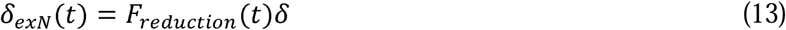

The F_rp_ represents the multiplier of potential root-produced auxin (*IAA_r,pot_*) to calculate the actual auxin produced in the root meristem in response to the external nitrate conditions. The F_reduction_ represents the multiplier of the combined rate for IAA reduction to calculate the rate of auxin reduction in the root meristem in response to the external nitrate conditions. The b_rp_ represents the coefficients of the relative root-produced IAA in response to external N concentration and b_reduction_ represents the coefficient of the relative IAA reduction rate in response to external N concentration. The δ represents a constant auxin efflux rate regardless of external N levels. Although external N concentration affects both auxin efflux and potential root auxin production in the model, the result in root total auxin remains robust across a range of relative strengths of these processes (Fig. S6).

### Model evaluation

To isolate the direct effects of the newly implemented auxin module and assess the robustness of its predicted effects, we first performed simulations with a fixed root biomass allocation (Supplementary Section 5.1). Then, to evaluate whether the model can capture major trends in root architectural responses across N treatment, we enabled all feedbacks in the plant model, leading to biomass allocation to the root being the result of biomass production through photosynthesis mediated by N uptake by the roots (‘*dynamic biomass allocation*’), and evaluated root length per 10 cm soil layer (Rootlength, m), shoot and root biomass and N uptake at all N levels with an experiment (see ‘*Experiment*’ below).

#### Evaluation scenario: dynamic biomass allocation

The simulation scene was set with 16 individual plants with 0.12 m of row distance and 0.025 m of plant distance and cloned 10 times on both x and y axis to construct a comparable density (343 plants/ m^2^) to the experiment. The simulations were run for 49 days with 10 replications across 4 N levels (LowN: 500, MedLowN:1000, MediumN: 2000, and HighN: 3000 µmol/L. From root length, the proportion of root length per layer (eq. 14) was derived as the fraction of total root length in the layer. The rest of the plant characteristics were derived as relative values as a fraction of maximum values across N levels (eq. 15).

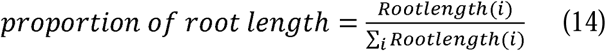

where i is an index for each soil layer.

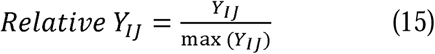

where Y_I_ represents shoot biomass, root biomass, root:shoot ratio and N uptake. J represents the different N levels.

#### Experiment

Data for model evaluation, was obtained from an experiment that was conducted from March 18^th^ to May 6^th^ 2020, at Wageningen University and Research, the Netherlands (Wang, 2025). Containers of size 70 cm × 90 cm × 40 cm were used as mesocosms. The containers were positioned beneath a transparent rain shelter in an outdoor environment. The average temperature over this period was 12.3 °C and the temperature sum was 602 °Cd using a base temperature of 0 °C.

The growth medium was sandy field soil mixed with river sand (1:1, v/v). The total N content of the soil before mixing was 14.6 mg/kg soil. 8.4g of 45% P_2_O_5_ and 5.6g of 30% K_2_O were applied uniformly. The soil depth was 36 cm with a dry bulk density of 1.39 g cm^-3^.

For each container, wheat (*Triticum aestivum* L. cv. Nobless) was planted in 6 rows with 12 cm between-row distance, and there were 36 plants within each row of 2.5 cm (343 plants/m^2^). A randomized complete block design with three replications was used. The NH4NO3 solution was used as N source for the four N treatments. The four N treatments were: LowN: 0g m^-2^, MedLowN: 2g m^-2^, MediumN: 4g m^-2^, and HighN: 6 g m^-2^. The N application was split into two equal parts, administered at 20 and 32 DAS.

Plant sampling was conducted on 35 days after sowing (DAS) and 49 DAS using the monolith method, lifting a soil block measuring 24 cm in length, 10 cm in width and 36 cm in depth. Sampling sites were in the middle rows to minimize the border effects. Aboveground biomass was collected and oven dried at 70 C for 48 hours. Shoot N was analyzed by using the Kjeldahl method. Roots were evenly divided into three parts based on their location in the sampled soil (12 cm each). After scanning the root, the software WinRhizo Pro (Regents Instruments Inc., Quebec City, Canada) was used to analyze root morphological characteristics such as root length to calculate the proportion of root length across three soil layers (12 cm each). Roots were oven-dried afterwards and the dry weight recorded.

### Using the model to analyse the role of root responses to N availability in plant N use

After having evaluated model functioning and output, we performed simulations to address the research question on the effects on root system architecture and plant N use of both systemic and local root responses to N availability. To do this, we first fixed root biomass allocation, which means there was no photosynthesis enabled and the biomass allocation was the proportion of the potential growth, to analyze only the direct effects of local and systemic responses to N without feedback effects on biomass accumulation and allocation. Afterwards, we enabled the full model again, enabling the feedback of N uptake due to the plastic responses on assimilate production and allocation throughout the plant. All environmental parameter values of all scenarios are listed in Table S4.

#### Scenario 1 - fixed biomass allocation to the roots

We ran the single plant simulations for 70 days until the root system fully developed with the systemic and local responses either both disabled, one of the responses enabled, or both enabled. To explore the effects of systemic and local root auxin responses on N uptake and avoid further root architectural changes due to changes in root biomass, the root biomass allocation was fixed as previously. We simulated 5 different N levels to create a large range of N availability in order to quantify the effect of plasticity: 10, 30, 300, 1000, and 3000 µmol/L. With the systemic responses disabled, the shoot-derived auxin values were set to the same values as those when plant N concentration would be high (0.0478g N/g DW). With the local responses disabled, no auxin export from root tips occurred, and the root-derived auxin was fixed to the value representing high N (3000 µmol/L). To account for variability in the model output caused by stochasticity in the model, such as the probability used in eq. 4 and eq. 7, the model was run 20 times for each treatment.

For each run, N uptake per plant and root length across the soil profile were recorded. The relative root length across the soil profile and N uptake per plant with one or both responses enabled in relation to root length/N uptake with all responses disabled, were calculated (eq. 16).

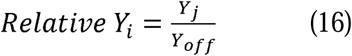

Yi can represent N uptake per plant, root length across the soil profile, total plant biomass, or plant N concentration. Y_off_ represents both responses disabled. Y_j_ can represent either or both responses enabled. Relative N uptake and root length were calculated to quantify auxin responses of root system architecture and their relationship with N uptake (Scenario 1).

Relative total biomass was calculated to evaluate the effects of different auxin responses on growth (Scenario 2). Relative plant nitrogen concentration was used as an integrative indicator of plant nitrogen status, accounting for biomass- and uptake-driven effects.

#### Scenario 2 – dynamic biomass allocation to the roots

To explore the relation between soil N, responses to N level and N uptake, we enabled the shoot part of the model and simulated light interception and N-dependent photosynthesis to drive biomass production. In this scenario, since the difference in total biomass among treatments is the output of interest rather than the absolute biomass, leaf senescence was not taken into consideration to reduce complexity caused by non-essential feedback for our research question. We ran simulations for 70 days with either the systemic or/and local responses either disabled or enabled, under 5 different N levels: 10, 30, 300, 1000 and 3000 µmol/L. Also, the model was run 20 times for each treatment combination. N uptake per plant, total plant biomass, root: shoot ratio, and plant N concentration were recorded. The relative total plant biomass, relative plant N concentrations, and relative N uptake with one or both responses enabled in relation to the same outputs with all responses disabled were calculated (eq. 16).

## Results

### Model evaluation

The effects of individual auxin-related parameters and processes on root length and root number across soil profile and effective auxin levels in the root meristem for high (3000 µmol/L) and low N (30 µmol/L) are shown in Fig. S6 to Fig. S9. Details have been described in the supplementary result section 5.1.

With dynamic biomass allocation enabled in the simulation, the aboveground biomass increased with increasing initial soil N from 500 to 1000 umol/L, but no difference for higher N levels (Fig. 2A). The root biomass and root:shoot ratio decreased with increased soil N (Fig. 2B and 2C), but still plant N uptake increased with increasing soil N (Fig. 2D). In the experiment, shoot biomass did not always clearly increase with N application which was in line with our modelling results (Fig. 2A). Also, root biomass and the root:shoot ratio reduced with increasing N application in line with model output (Fig. 2B, C). N uptake increased with increasing N applications, as in the model output (Fig. 2D). More roots were distributed in the top soil layers at high N than at low N, while more roots were distributed in the deeper soil layers at low N than at high N (Fig. 3A). We observed a similar trend in the proportion of root length per soil layer in the mesocosm experiment (Fig. 3B).

**Fig. 2.**
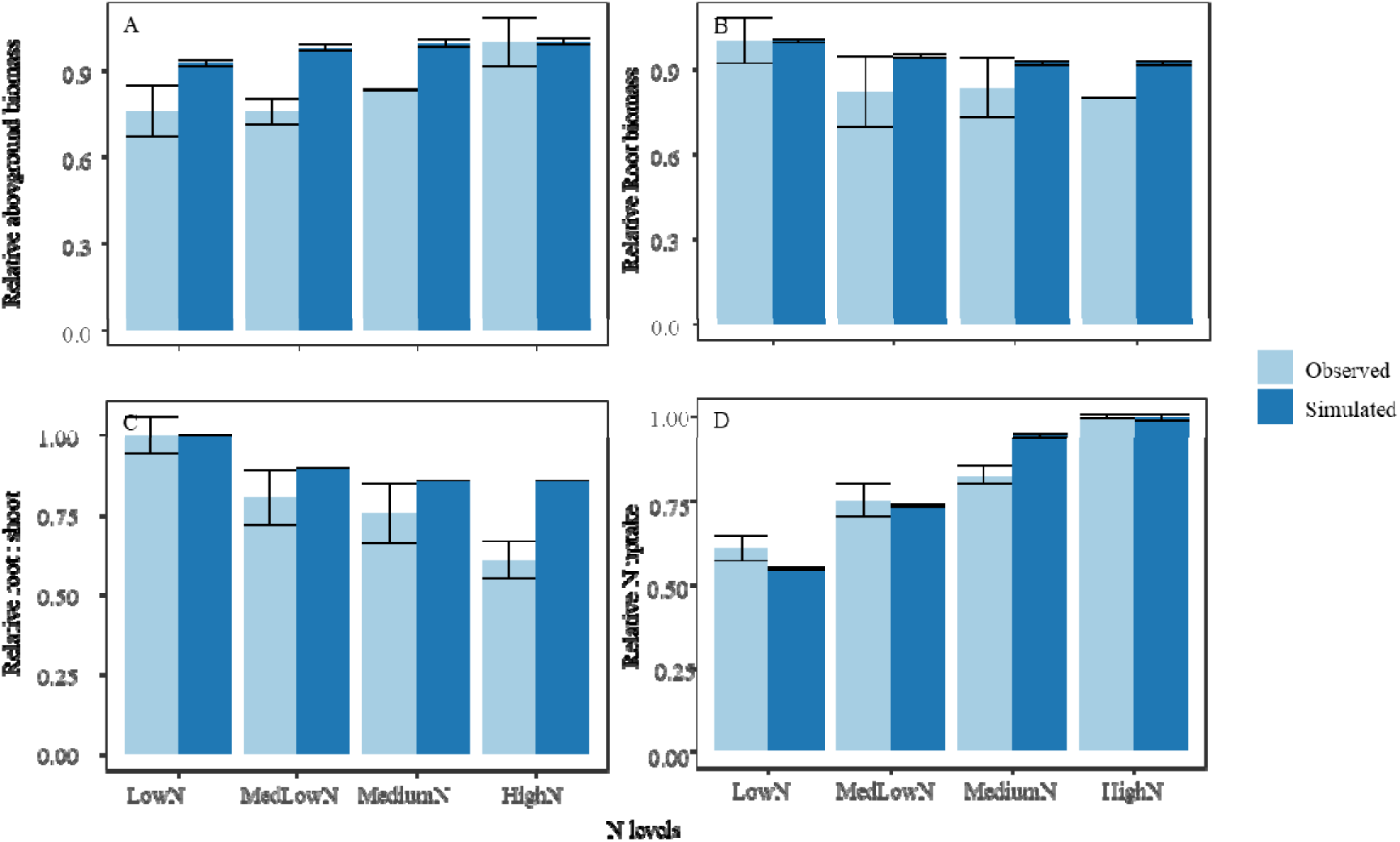
Simulated vs observed plant characteristics across multi-N levels at planting density of 343 plants/m^2^. The LowN, MedLowN, MediumN and HighN represent total N levels in simulation: 500, 1000, 2000, and 3000 µmol/L. In the experiment, the LowN, MedLowN, MediumN and HighN represent applied N in the soil: 0, 2, 4, and 6 g/m2. The simulations begin at the 77^th^ day of the year and run for 49 days after sowing. Values are mean ± SE (n_sim_=10, n_obs_=3). The absolute values were shown in Supplementary Fig. S11 and S12.

**Fig. 3.**
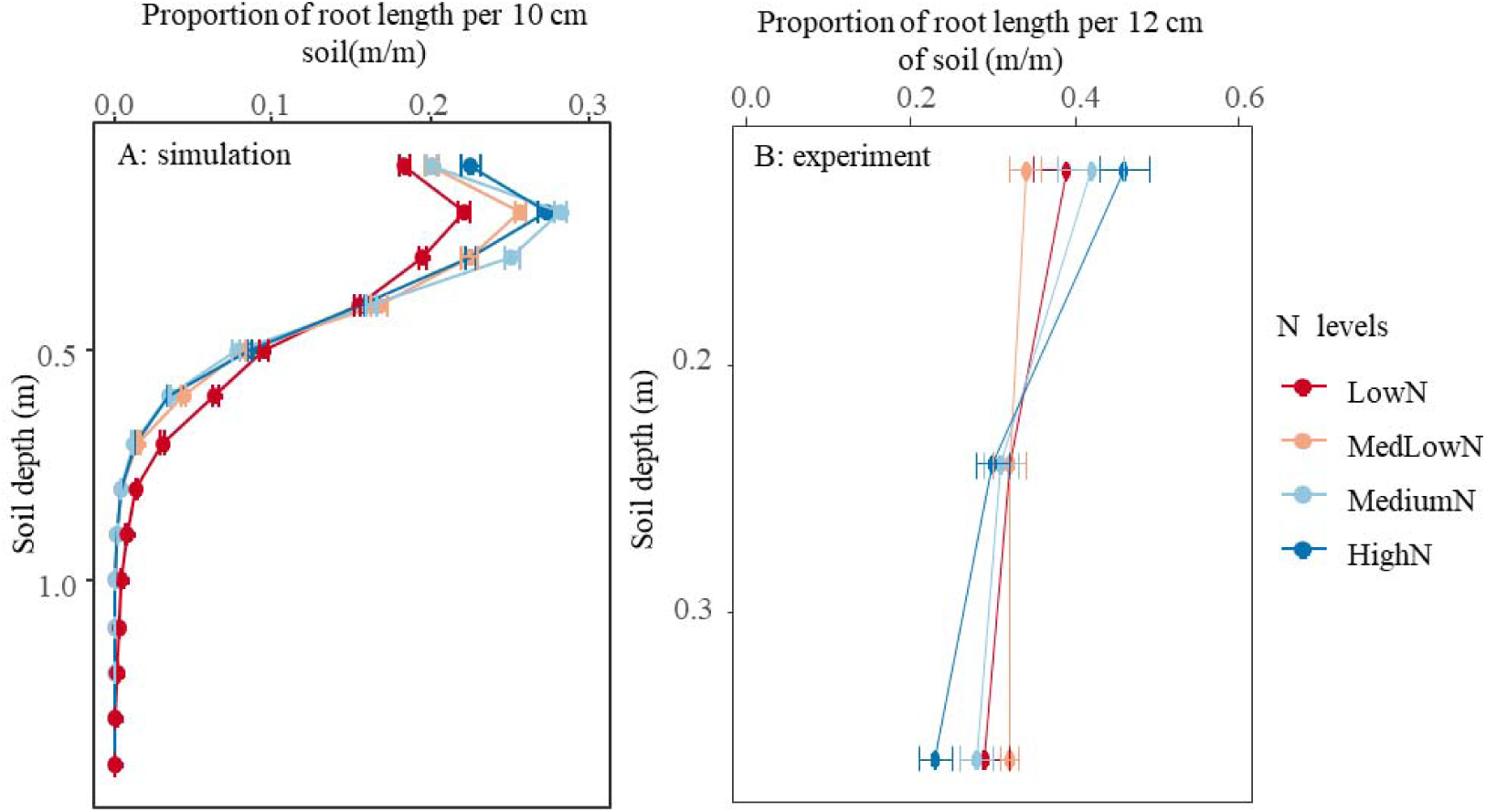
Simulated (A) vs Observed (B) proportion of root length (m/m) across soil layers. The proportion of root length is the fraction between root length in the individual soil layer and total root length. The LowN, MedLowN, MediumN and HighN represent total N levels in simulation: 500, 1000, 2000, and 3000 µmol/L. In the experiment, the LowN, MedLowN, MediumN and HighN represent applied N in the soil: 0, 2, 4, and 6 g/m^2^. Values are mean ± SEs (n_obs_=3, n_sim_=10).

### Exploring the effect of local/systemic responses

#### Scenario 1 - fixed biomass allocation to the roots

When the plant faced low N (10 and 30 µmol/L), disabling the systemic response to N increased the N uptake compared to both responses enabled (Fig. 4). Both these situations resulted in a relative N uptake higher than 1.0, meaning N uptake in those cases exceeded N-uptake in the absence of responses. In contrast, disabling the local response decreased the relative plant N uptake to below 1.0, also at moderate N (Fig. 4). When N levels were high enough (1000 and 3000 µmol/L), no differences in N uptake were found between responses (Fig. 4). There were continuously increasing trends of relative N uptake at 3000 µmol/L especially when systemic response enabled (Fig. 4).

**Fig. 4.**
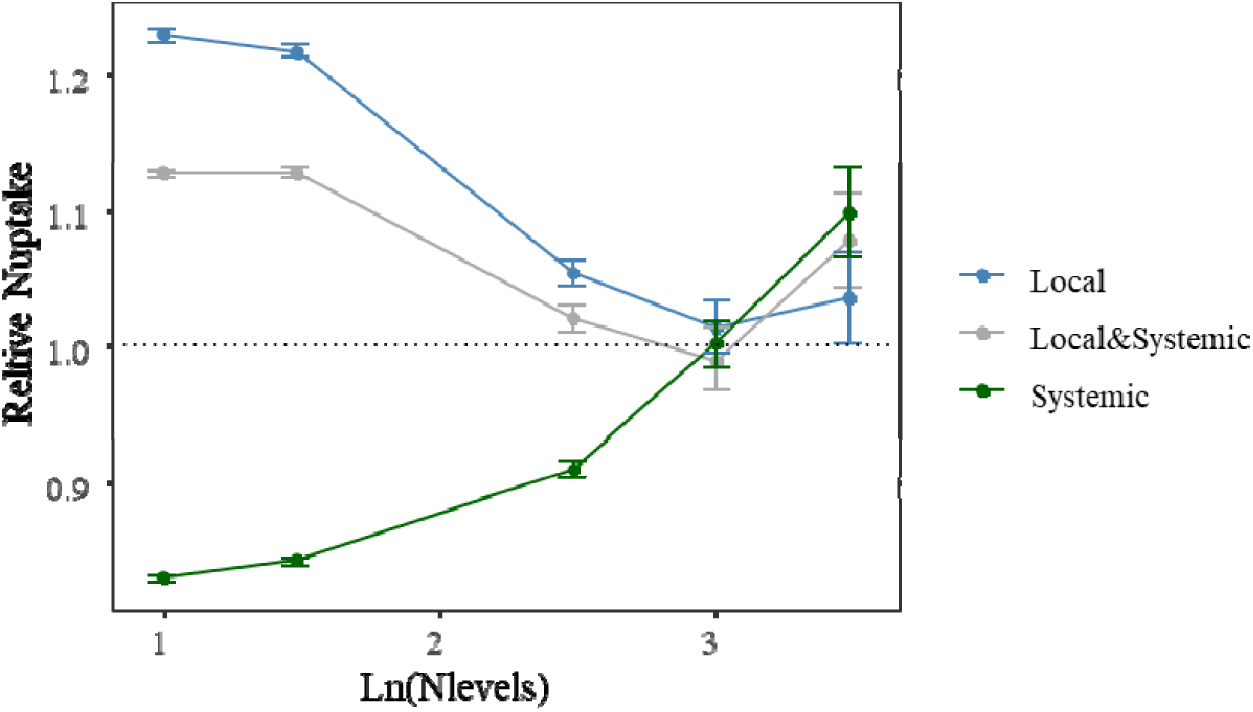
Simulated N uptake for different response types, relative to N uptake without either response, in the fixed-biomass-allocation scenario. The simulations were run for 70 days. The dotted line represents a relative N uptake of 1. ‘Local’, ‘Local & Systemic’, and ‘Systemic’ indicate which responses to N were enabled in simulation. The x-axis represents the natural logarithm of initial soil N at 10, 30, 300, 1000, and 3000 µmol/L. The values represent mean ± SEs (n=20).

To further explore why there were differences in N uptake (Fig. 4) even though root biomass allocation had been fixed, the relative root length in the soil profile was analyzed (Fig. 5).

**Fig. 5.**
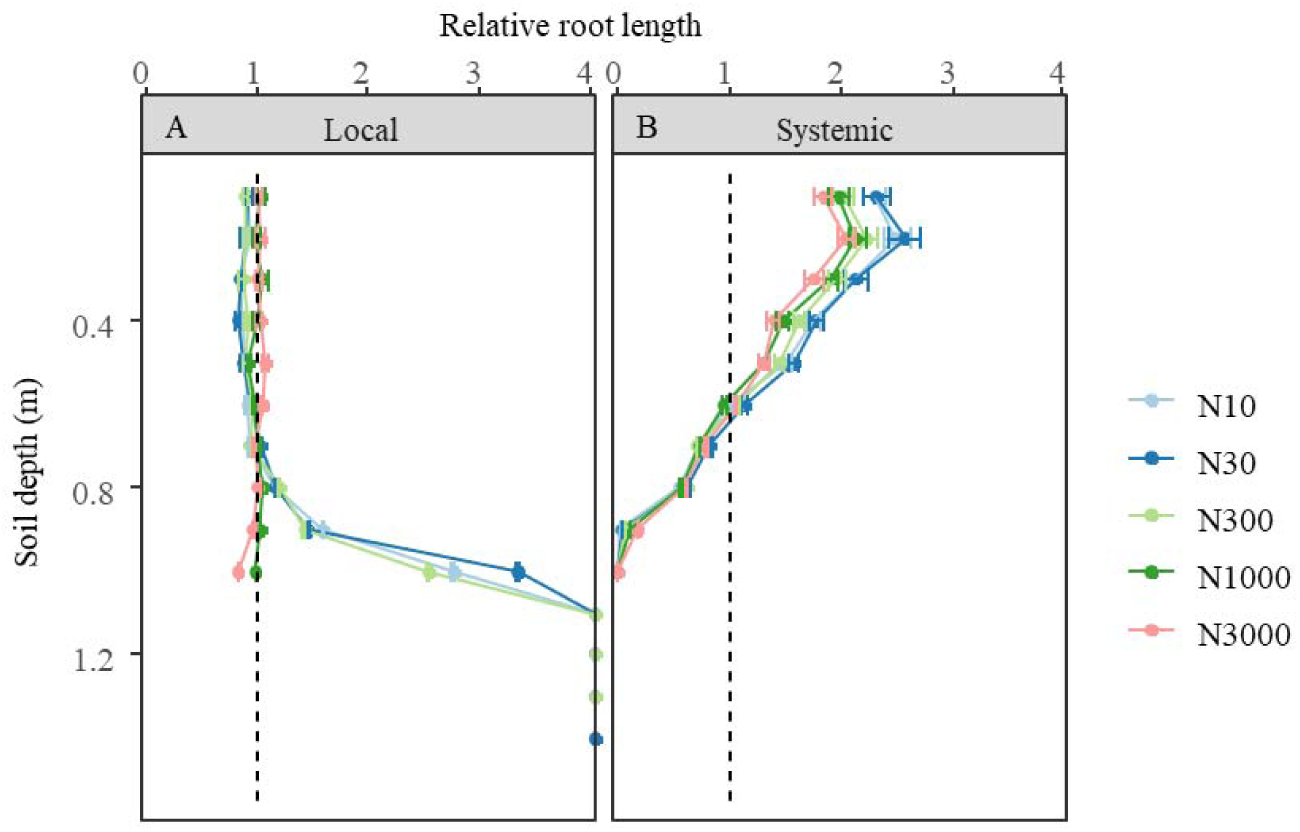
Simulated relative root length across all N levels calculated as the ratio of root length between with local (A) or with systemic (B) responses only and without any of the responses. N10, N30, N300, N1000, and N3000 represent the initial soil N at 10, 30, 300, 1000, and 3000 µmol/L. The dashed line represents a relative root length within each 10 cm soil depth of 1. The values represent mean ± SEs (n=20).

When only enabling the local auxin responses, more roots were distributed in the deeper soil profile than the roots with disabled auxin responses under low N, which enhanced foraging extra N in deeper soil (Fig. 5A). When only enabling the systemic auxin responses, more roots were distributed in the upper soil profile than the roots with disabled auxin responses (Fig. 5B). This also indicates that the continuous increase in relative N uptake under high N (Fig. 4), when the systemic response is enabled, is driven by dynamic changes in the probability of lateral root emergence (Fig. 5B). These dynamic changes cannot be captured when both responses are disabled.

#### Scenario 2- dynamic biomass allocation to the roots

When running the simulations with the plant C and N sink-source feedbacks enabled, less biomass with only the systemic response enabled was found compared with only the local response enabled under low N (Fig. 6). At higher N levels, the difference between enabling local or systemic responses became negligible (Fig. 6). The relative N concentrations with dynamic biomass allocation have less variation across the four response types than with fixed carbon allocation under low N (Fig. 7A and 7B) while no difference was found in variation of relative N concentration between dynamic and fixed biomass allocation for the high N.

**Fig. 6.**
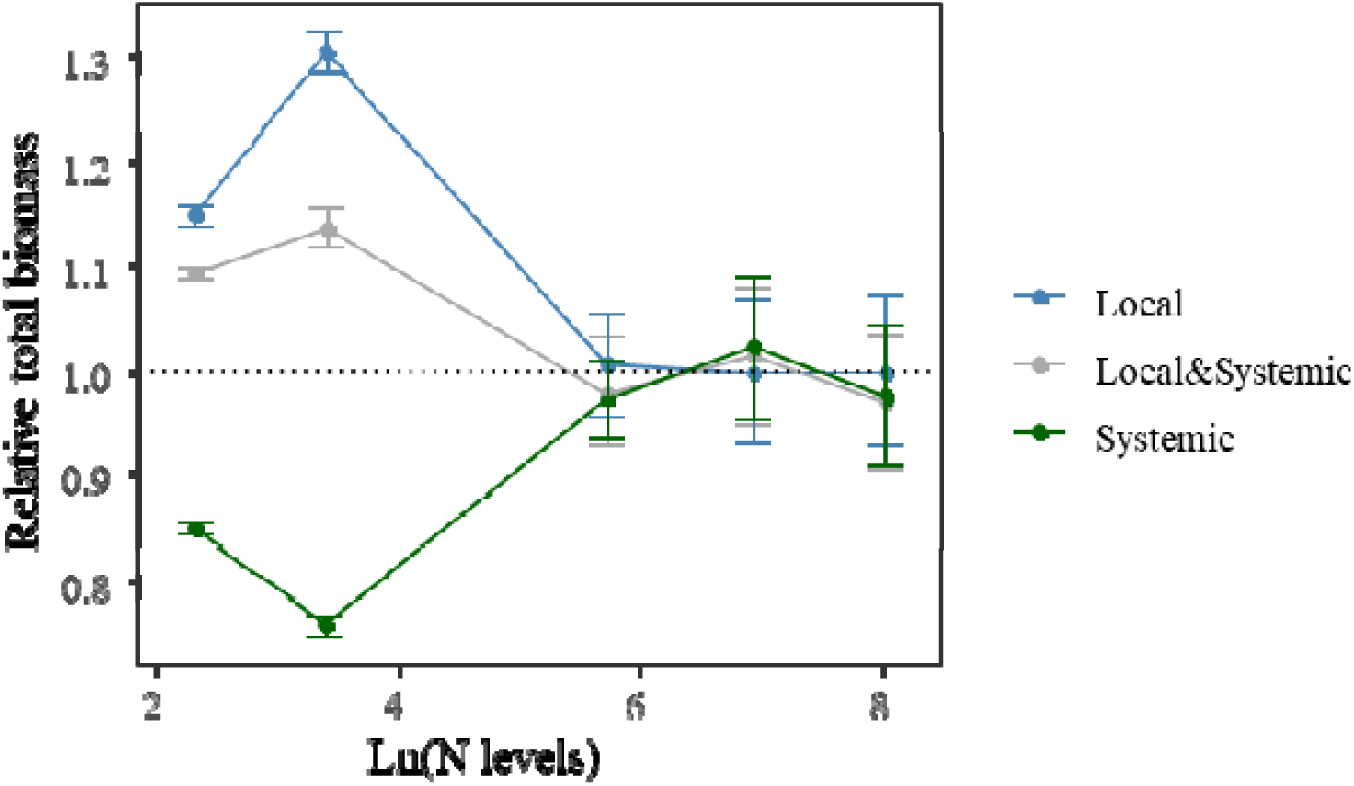
Simulated total biomass for different response types, relative to total biomass without either response, in the dynamic-biomass-allocation scenario. The simulations were run for 70 days. The dotted line represents a relative N uptake of 1. ‘Local’, ‘Local & Systemic’, and ‘Systemic’ indicate which responses to N were enabled in the simulation. The x-axis represents the natural logarithm of initial soil N at 10, 30, 300, 1000, and 3000 µmol/L. The values represent mean ± SEs (n=20).

**Fig. 7.**
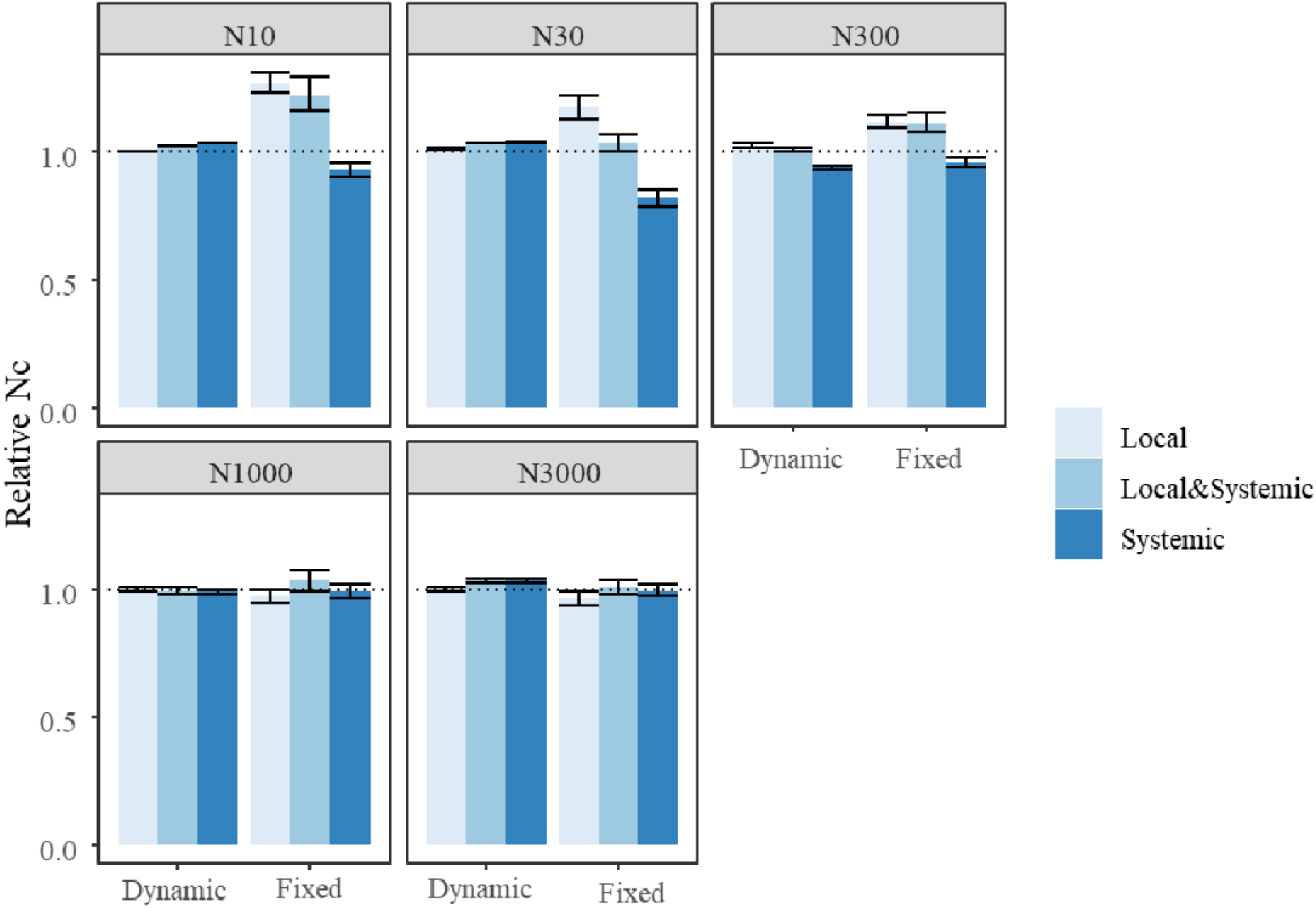
Simulated plant N concentration for different response types, relative to plant N concentration without either response. The simulations were run for 70 days. The “Dynamic” and “Fixed” indicate dynamic and fixed biomass allocation. The dotted line represents a relative N uptake of 1. “Local”, “Local & Systemic”, and “Systemic” indicate which responses to N were enabled in the simulation. N10, N30, N300, N1000, and N3000 represent the initial soil N at 10, 30, 300, 1000, and 3000 µmol/L. The values represent mean ± SEs (n=20).

## Discussion

In this study, we explored and quantified the effects of auxin-driven local and systemic responses in root architectural development to plant N availability, and the consequences for N uptake and plant growth. Simulations were first run under high and low N treatments for 20 days while changing values of auxin-related parameters and processes, to evaluate the extent to which the auxin responses played roles in root functioning and architectural development (Fig. S7; Fig. S8). Changing the levels of shoot- or root-derived auxin (Fig. S7) did not alter the major trends in the simulated responses to high and low N, indicating that the conclusions are robust to variation in auxin levels. Combined with the sensitivity analyses presented in Supplementary Fig. S7, Fig. S8 and Fig. S9, Fig. 5 shows that under external low N, when only local responses were enabled, reduced effective auxin in the root tip resulted in increasing the potential elongation rate of axial roots, resulting in deeper rooting compared with high N. In contrast, when only systemic responses were enabled, low plant N increased the shoot-derived auxin in the root tips, thereby increasing the probability of lateral root emergence during early growth stages. Consequently, more lateral roots developed in the upper soil layers than under high N. Our simulation results (Fig. S7 and Fig. S8) can reproduce the observations that shoot-derived auxin plays an essential role mainly in lateral root emergence, whereas both root and shoot-derived auxin play a role in lateral root initiation (Du et al., 2018; Hu et al., 2021).

When running simulations with dynamic biomass allocation enabled, the trends of important model outputs align with those we observed in the experiment (Fig. 2, 3). The simulated trends in root:shoot ratio, root biomass, shoot biomass, and N uptake were in line with observations in additional studies as well, across many crop species, including wheat, in both pot and field experiments (Kamiji et al., 2014; Lopez et al., 2023). In addition, various studies have shown that a higher root length is found in the top soil under high N compared with under low N, while roots are more distributed towards the bottom soil under low N than under high N (Chen et al., 2014; Lynch, 2013; Lynch et al., 2024). The results of the model evaluation lead us to conclude that the model is a valid tool for exploring the effects of local and systemic responses to N on root architecture and plant growth.

### Tolerance vs avoidance under low N conditions

In our modeling, we implemented two different strategies for survival under low N conditions, through local and systemic auxin-related responses in the root system to N (Fig. 1).

The local responses led to a direct increase in root N uptake under low N by increasing root length distribution in the deep soil layers (Fig. 4 and Fig. 5A). Under moderately low N, plants can upregulate auxin local synthesis genes, which can promote elongation of the lateral and axial roots (Gruber et al., 2013; Jia et al., 2021; Yu et al., 2014) leading to improved N foraging capacity (Lynch, 2013). While at an extremely low N condition, the benefit of local responses may not be sustained (Fig. S14; Wang et al., 2023).

The systemic responses, on the other hand, led to more roots in the topsoil layer, which could lead to a shallow root system architecture and less N uptake under low N (Fig. 6 and Fig. 7B). Such shallow root architecture is not ideal for low N situations, due to less root forage capacity for water or N, which can easily leach out of the system (Lynch, 2013; Saengwilai et al., 2014a, 2014b; York et al., 2015). Under low N conditions, reduced N uptake can result in less biomass accumulation (Fig. 6), reducing the N demand. As a result, both local and systemic auxin-related responses can maintain plant N concentration requirements under low N (Fig. 7).

In theory, local responses to N could belong to a stress-avoidance strategy by forming a deep root system to forage for extra N and complement surface N-deficient conditions (Berger et al., 2016; Volaire, 2018). This type of strategy enhances plant competitiveness and is useful when a plant is facing moderate stress (Berger et al., 2016; Grime, 1974; Qi et al., 2018). The systemic responses to N could belong to a stress-tolerance strategy aimed at reducing plant size to survive under stress. The stress-tolerance strategy is considered to have low competitiveness, but can assist plants in overcoming and surviving strong stress conditions (Qi et al., 2018). The local and systemic responses exhibit a trade-off in plant N uptake, which is also in line with the trade-offs observed between avoidance and tolerance for other abiotic and biotic forms of stress (Henry et al., 1997; Krimmel et al., 2016; Puijalon et al., 2011; Vesk, 2006).

By implementing responses in auxin biosynthesis, transportation, and signaling to local and systemic N (Fig. 1), our model captured the trade-off between avoidance and tolerance in low N situations. The coexistence of both responses in the plant simultaneously enables it to cope with both moderate and severe N stress. Under moderately low N stress, local responses could play a major role in assisting plants to overcome the stress by enhancing their forage ability.

This phenomenon has been observed in several studies (Jia et al., 2021, 2022; Jia & von Wirén, 2020). Interestingly, towards more severe N stress, the systemic signalling-related genes, such as the root-to-shoot mobile peptide hormone, C-TERMINALLY ENCODED PEPTIDE (CEP), are upregulated under N-starvation (Ohkubo et al., 2017). This fine-tuning system between systemic and local responses could lead the plant to react precisely and efficiently to survive across a wide range of N conditions.

Genotype selection under either moderate or extremely low N environments can lead to enhancing either avoidance or tolerance characteristics separately, and may result in unexpected negative effects. The small size of a tolerant plant may result in reduced plant productivity, as smaller photosynthetic organs may lead to the assimilation of less carbon; therefore, the potential final yield would be lower (Michaletz et al., 2014). On the other hand, plants with strong avoidance responses may increase competition among crop plants for resources. In our results, with strong avoidance (i.e., a strong local auxin response), a higher fraction of biomass supports root growth, especially at the early growth stage (Fig. S14), which could be an additional constraint on biomass formation for reproductive organs. Rather than focus on one or a few specific environments, genotype selection for plasticity also needs to be conducted with a wide range of environments to reduce the risk of maladaptation to alternative environments resulting from excessive selection for particular plasticity.

### Applying FSP models to explore complex interactions

In this study, we expanded established FSP modelling concepts by implementing an auxin module in the root tips taking care of responses to N. We included the auxin signalling in the model and linked them to potential root elongation rate and lateral root development. Such expansions provide opportunities to explore the complex interactions among plant signals, architecture, growth, and development. In this study, we applied the model to explore the effects of auxin-related systemic and local responses on plant resilience under low N conditions (Fig. 4 to 7). These effects usually interact highly with each other and can hardly be studied individually, even when using mutants (Tonsor et al., 2005; Ye et al., 2025). This FSP modelling approach can disentangle the highly interacting effects at any developmental stage, which can help to uncover the effects that may be hidden in real plants. This use of FSP modelling has been demonstrated by Prusinkiewicz et al. (2009) to understand the control of bud break and branch development based on auxin signalling mechanisms. In a previous study, we disentangled the effects of active plasticity (i.e. changes in plant traits due to changes in relative sink strength or development) and passive plasticity (i.e. changes in plant traits due to resource limitation) of the root system through this type of modelling approach to study how active plasticity influences resource capture (Lu et al., 2024a). Such modelling approaches can provide new insight into understanding the complex interactions within a plant and between plants and the microenvironment.

Auxin has been widely modeled at the cellular level (Allen & Ptashnyk, 2020; Grieneisen et al., 2007; van den Berg et al., 2021). Such models can well interpret auxin distribution within the plant tissue and interactions between auxin transportation and growth of root tips. Due to the complexity of auxin mechanisms within the plant, cell-level tissue models can hardly be directly up-scaled to explore how auxin influences plant functions at the whole-plant level or even at the plot level, which is important for our understanding of the functioning of crops or other vegetations. Based on this, our FSP modelling approach with a simplified auxin module could be an appropriate tool to study auxin functions at the whole-plant scale, studying inter- and intraspecific competition for resources in natural and crop settings.

There are still limitations to our modelling approaches. The explicit or accurate parameterization for every relevant process is impossible due to a lack of knowledge or data. For instance, the threshold value (*V_auxin_*) to account for the cell function disruption under an extremely low N situation turns out to be sensitive to the local response and to be species-specific, requiring further study (Fig. S15; Sun et al., 2020; Wang et al., 2023). Therefore, we decided not to include this constraint in the current model explicitly. Our current study only focused on a limited number of environmental factors driving plant growth and development (light, temperature, and nitrogen), but environmental stress is always caused by multiple environmental factors. To understand how environmental stress influences plant growth, other environmental factors, such as drought, can be further considered. In the current model, we include the feedback of auxin in the root tips. Further including plant signal processes in the whole plant model (Evers et al., 2011; Runions et al., 2017) can provide an extra understanding of the complex interaction among plant signaling, function, structure, and environments.

## Conclusion

Through 3D plant modelling, we show that local auxin responses directly enhance N uptake by distributing more roots towards the deep soil under low N conditions, while systemic auxin responses reduce N uptake by distributing more roots near the top soil layers. These two different strategies can both contribute to overcoming low N stress. The local auxin responses to N represent stress avoidance by enhancing root growth, which could form extra roots and explore the extra space. The systemic auxin responses represent stress tolerance by reducing plant size, which reduces the demand for N. There is a trade-off in N uptake between the two strategies, and the trade-off can be explained through C or N allocation and demand to different plant organs. This modelling study further illustrated that this modelling approach is a powerful tool to disentangle the complex interactions in real plants to understand the effects and interactions between mechanisms behind strategies to cope with N deficiency.

## Acknowledgement

The authors are grateful to Dr. Thea van den Berg and Prof. Lixing Yuan for providing insightful thoughts to improve the manuscript. The study was financially supported by the European Union (EU horizon project IntercropVALUES, grant agreement No 101081973).

## Competing interests

The authors declare no conflicts of interest.

## Author contributions

JL and JBE designed the research; JW performed the experiment. JL, JBE and AM developed the model and analysed data; JL wrote the manuscript. JL, JBE, AM and JW revised the manuscript and contributed to the finalization of the manuscript.

## Data availability

The model code will be made publicly available in an open-source repository upon publication.

## Supporting Information

### Auxin module parameterization

Fig. S1: Parameterization for probability of lateral root outgrowth as a function of rootward auxin concentration.

Fig. S2: Parameterization for the estimated factor of potential axial root elongation rate as a function of auxin concentration.

Fig. S3: Parameterization for the estimated probability of initiation of the lateral root (Pinit) as a function of auxin concentration.

Fig. S4: Parameterization for the estimated factor of IAA concentration from the shoot as a function of total plant N concentration.

Fig. S5: Parameterization for the multipliers of potential root-produced auxin and auxin reduction rate in the root meristem at the given external N.

### Model evaluation

Fig. S6: Effective total IAA vs days in response to consistent external N and internal plant N concentration.

Fig. S7: Simulated root length and lateral root number on the second axial root across the soil profile.

Fig. S8: Simulated changes in root length and lateral root number across the soil profile.

Fig. S9: Effective auxin in axial root with shoot/root auxin levels under high N (3000 µmol/L) and low N (30µmol/L) conditions.

Fig. S10: Effective auxin in the axial root of disabled auxin-related processes under high N (3000 µmol/L) and low N (30 µmol/L).

Fig. S11: Simulated actual axial root elongation rate vs axial root effective auxin under high N (HN, 3000 µmol/L) and low N (LN, 30 µmol/L).

Fig. S12: Simulated plant characteristics (absolute values) across multi-N levels. Fig. S13: Observed plant characteristics (absolute values) across multi-N levels.

### Exploring the effect of local/systemic responses

Fig. S14 Simulated root-to-shoot ratio over days across N levels. In the simulations, whole plant feedback and photosynthesis were enabled.

Fig. S15 Simulated total biomass for different response types, relative to total biomass without either response.

## Supplementary Tables

Table S1: Definitions of auxin-related variables

Table S2: Auxin-related parameter values

Table S3: Plant level parameter values

Table S4: Environmental parameter values

## Notes

### Competing Interest Statement

The authors have declared no competing interest.

