## Supplemental Data 1 for "Emergence of tolerance and avoidance strategies from local and systemic responses to nitrogen: insights from modelling of auxin-mediated root plasticity"

1. **Auxin module**
   1.1. The function of auxin in root meristem

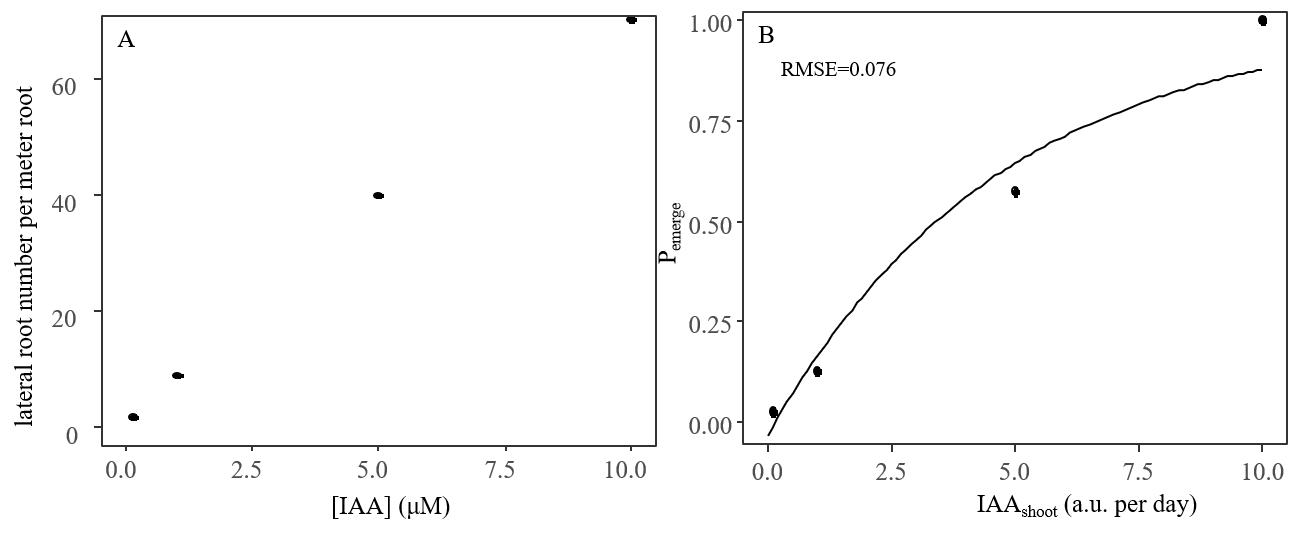

Figure S1. Lateral root numbers per unit root length as a function of auxin concentration at root-shoot junction (A) and probability of lateral root outgrowth as a function of rootward auxin concentration (B). Dots in panel A are the measured lateral root number per meter root from Reed et al. (1998). Dots in panel B are the normalized lateral root number per meter root used to fit the model (eq. 4 in the main text). The unit a.u. here stands for auxin unit.

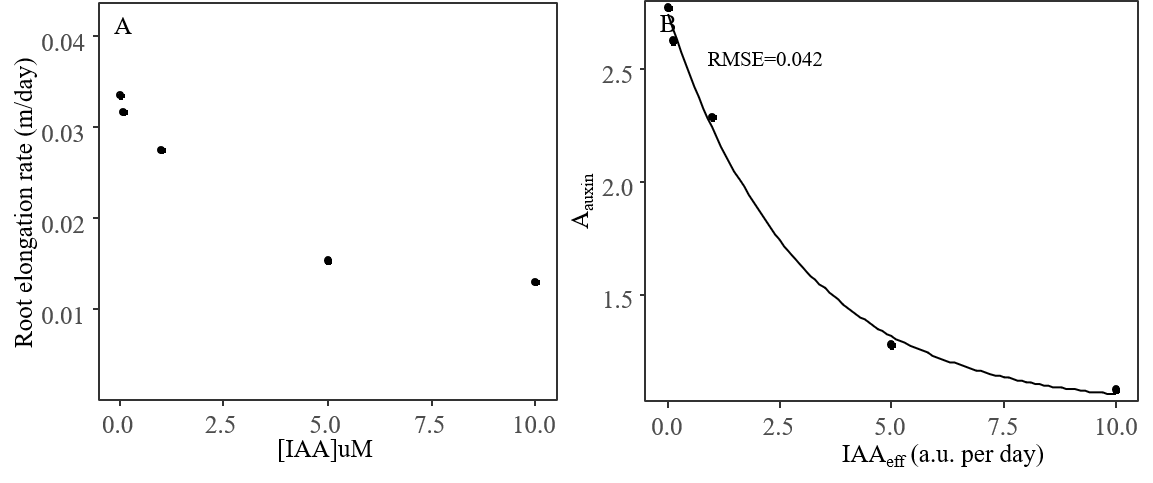

Figure S2. Primary root elongation rate as a function of auxin concentration (A) and estimated factor of potential axial root elongation rate as a function of auxin concentration (B). Dots in panel A are the measured data from Reed et al. (1998). Dots in panel B are the normalized lateral root elongation rate used to fit the relationship (eq. 5 in the main text) used in the model. The unit a.u. here stands for auxin unit.

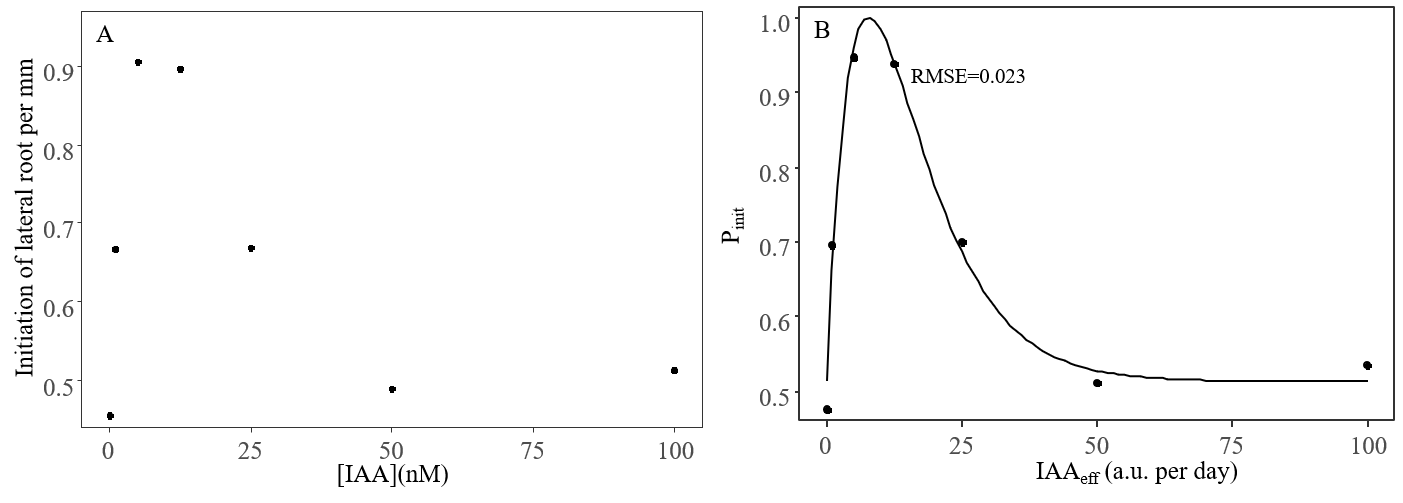

Figure S3 Initiation of lateral root per mm as a function of auxin concentration (A) and estimated probability of initiation of the lateral root (Pinit) as a function of auxin concentration (B). Dots in panel A are the measured data from Ivanchenko et al. (2010). Dots in panel B are the normalized lateral root primordia per meter root used to fit the probability model (eq.7 in the main text). The unit a.u. here stands for auxin unit.

1.2. Auxin translocation in response to internal plant nitrate

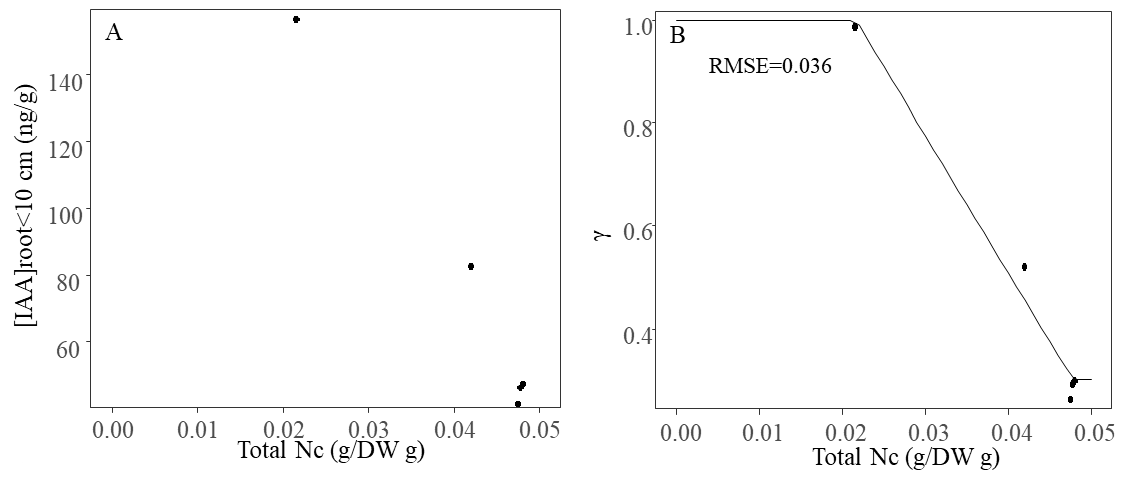

Figure S4 IAA concentration at root tip (<10cm) as a function of total plant N concentration (A) and estimated factor of IAA concentration from shoot as a function of total plant N concentration (B). Dots in panel A are the measured data from Tian et al. (2008). Dots in panel B are the normalized [IAA] translocated from the shoot used to fit the model (eq. 8 in the main text).

1.3. Auxin biosynthesis and translocation in response to external nitrate

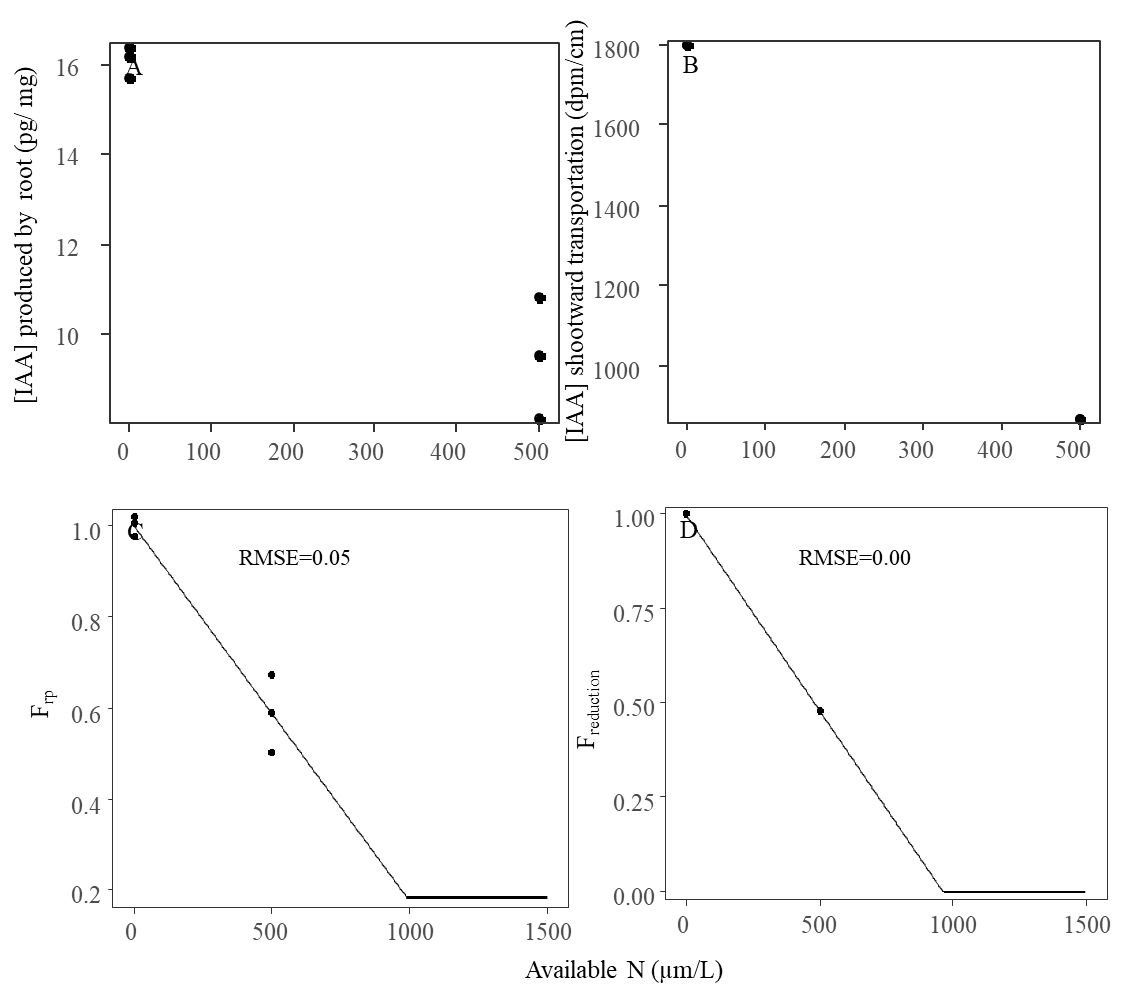

Figure S5 IAA concentration produced by root vs N application in a split-root system (A) and shootward transportation of IAA from root tip (measured at 3 cm to 6 cm from root tip) vs N application in a split-root system (B). Panel C (eq. 10 in the main text) represents the [IAA] produced by the root at the given N level and Panel D (eq. 12 in the main text) represents the multiplier of the reduction rate of [IAA] from the root tip at given external N. The Dots in Panel C and D are the normalized [IAA] measured in Liu et al. (2010) to fit the model.

2. **Intermediate step to calculate total auxin in one root meristem**

To assess the robustness of the interaction between auxin efflux and root auxin production in response to external N concentration, we implemented the auxin module for a single axial root meristem separately in R (version 4.3.2). For each external N level, internal plant N concentration, potential shoot auxin rate, and potential root auxin rate were held constant. This intermediate analysis isolated the combined effects of external N on root auxin production and auxin efflux.

| 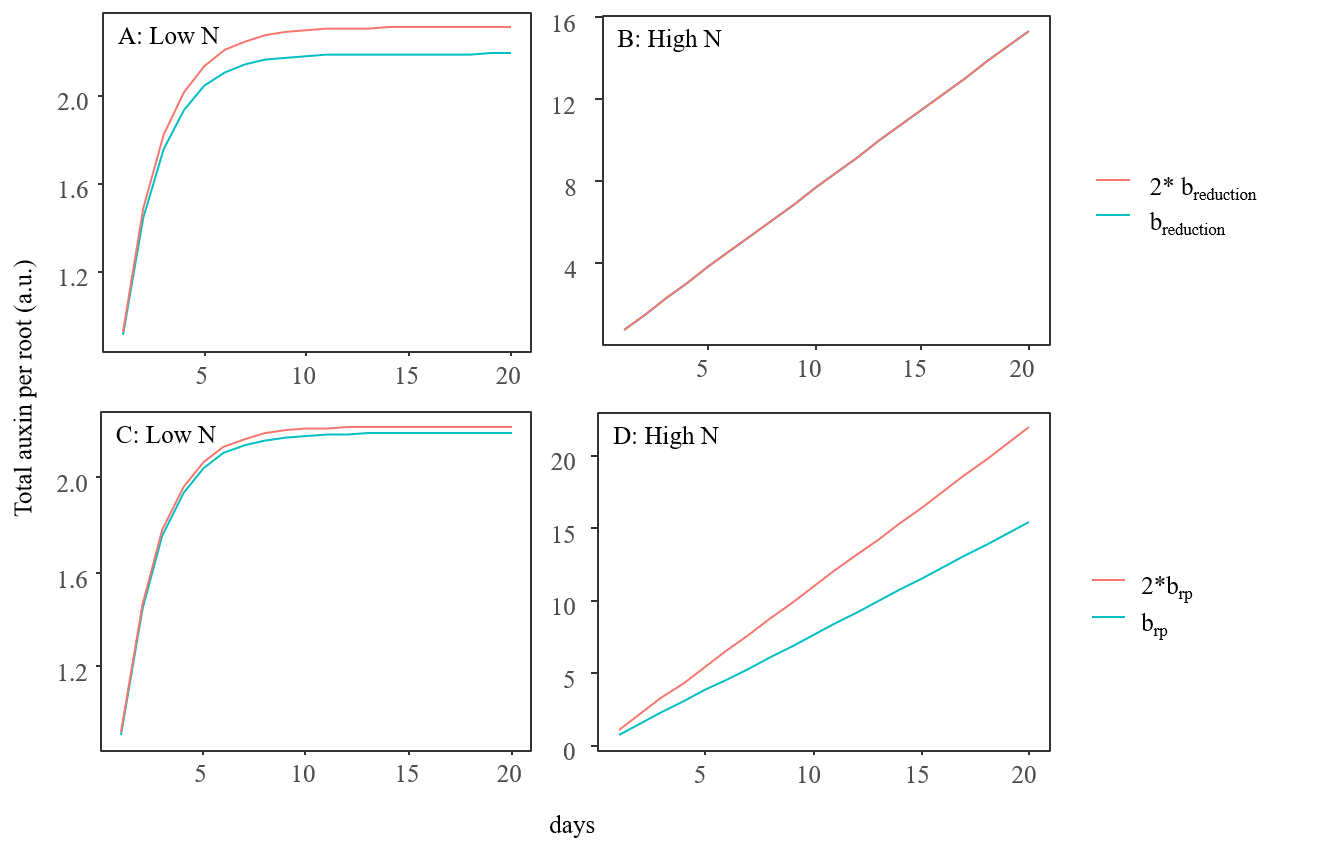 |
| --- |
| Figure S6 Effective total IAA vs days in response to consistent external N (High N: 3000 umol/L and Low N: 30 umol/L) and internal plant N concentration. The simulated days were 20 days. The b_reduction_ represents the coefficient of the relative IAA reduction rate in response to external N concentration in eq. 12. The brp represents the coefficients of the relative root-produced IAA in response to external N concentration in eq. 10. |

3. **FSP model description**

Our wheat FSP model represents the development and growth of both root and shoot in 3D and can be driven by light, temperature, and soil N (de Vries et al., 2021; Evers et al., 2010; Evers et al., 2007; Lu, et al., 2024b). The simulation runs on a daily basis. The plant and environment parameter values can be found in Table 3 and Table 4.

3.1. Carbon sinks

The model considers individual leaves, internodes, and grains as individual carbon sinks. The root system is considered one sink. Assimilated carbon is allocated to plant organs based on the individual organ sink strength (ss, g), which is defined as a sigmoid function of the organ potential dry mass (w_max_, g) and growth duration (t_e_, ^o^Cd) (eq.S.1, S.2; Evers et al., 2010).

$maxsink(i)=\frac{2t_{e}-t_{m}}{t_{e}\left( t_{e}-t_{m} \right)}\left( \frac{t_{m}}{t_{e}} \right)^{\frac{t_{m}}{t_{e}-t_{m}}}w_{max}(i)$	(S.1)

$ss\left( i,t \right)=maxsink(\frac{\left( t_{e}-t \right)t}{(1-r_{g}){(t_{m})}^{2}})$	(S.2)

where *i* represents the organ type (i.e. leaf, internode, grain and root system) and *t_m_* is assumed equal to *te*/2, which reduces the number of parameters but still well represents wheat growth. The parameter r_g_ is the fraction of potential substrate lost to growth respiration (Evers et al., 2010).

Even though the root system is treated as one sink for assimilates, it is further separated into axial roots, first-order lateral roots, and higher order lateral roots. The carbon sink strength of each root can be calculated as a function of potential root elongation rate (EL_auxin_, eq.4 in the main text), root diameter (d, m), and root tissue density (RTD, g/m^3^) (eq. S.3). Assimilates allocated to the root system are divided over the individual roots based on these root sink strength values.

${ss}_{root}\left( i,t \right)={EL}_{auxin}\left( t \right)d\pi{(\frac{d}{2})}^{2}RTD$	(S.3)

where *i* represents the individual roots, including axial, first-order lateral, and higher order lateral roots. To reduce computational complexity, higher-order lateral roots are only treated as sink for assimilates that contribute to N uptake from soil but their 3D structures are not simulated explicitly.

The functional equilibrium theory (Shipley and Meziane, 2002), states that, when exposed to limiting resources, plants invest extra carbon to those organs relevant to acquiring the limiting resource. This concept has been adopted in the model as a root-to-leaf partitioning coefficient (Λ_root:shoot_) in response to plant N concentration (Nc, eq. S.4, modified from Lu et al., 2024a). This coefficient modulates the root sink strength.

$$\Lambda_{root:shoot}(t)=\left\{ \begin{aligned} a_{r}e^{{-b}_{r}Ncmin}+1, Nc\left( t \right)<Ncmin \\ a_{r}e^{{-b}_{r}Nc\left( t \right)}+1, Nc\left( t \right)\geq Ncmin \end{aligned} \right. (S. 4)$$

The a_r_ and b_r_ are the empirical coefficients. Ncmin represents the minimum plant N concentration.

3.2. Carbon source

All leaves and internodes perform photosynthesis as a function of organ light capture and nitrogen content. An asymptotic exponential relationship was used to estimate photosynthesis rate (eq. S.5). This relationship is widely used in different crop models, such as e.g. SUCROS (Archontoulis and Miguez, 2015):

$An=Amax(t)\times\left( 1-e^{\frac{-\alpha\times PAR}{Amax}} \right)-Rd$	(S.5)

where Amax (µmol CO_2_/(m^2^·s)) represents the maximum photosynthetic rate, α represents the initial slope of the curve at low irradiance levels, Rd (µmol CO_2_/(m^2^ s)) is the dark respiration rate, and PAR (µmol/(m2 · s)) represents photosynthetically active radiation. Amax is dependent on specific leaf nitrogen (SLN, g/m^2^):

$Amax\left( t \right)=\lambda*(\frac{2}{\left( 1+e^{\left( -a*(SLN-0.18 \right)} \right)}-1)$	(S.6)

where a and λ are empirical coefficients. Since internodes also contain chloroplasts and can perform photosynthesis, SLN also includes the photosynthetic nitrogen of the internode

The actual growth of each organ is co-determined by both carbon sink and source strengths (eq. S.7a). The actual growth of each root is co-determined by each root carbon sink (eq. S.3) and the actual assimilates allocated to the root system (eq. S.7b).

$growth\left( i,t \right)=min(sinkstrength\left( i,t \right), assimilates\left( t \right)\frac{sinkstrength\left( i,t \right)}{\sum_{i} sinkstrength\left( i,t \right)})$	(S.7a)

$growth\left( j,t \right)=\min\left( rootsinkstrength\left( j,t \right), growth\left( root,t \right)*\frac{rootsinkstrength}{\sum_{j} rootsinkstrength\left( j,t \right)} \right)$ (S.7b)

where *i* represents the leaf, internode, grain and root and *j* represents the individual axial root, first-order lateral root and higher-order lateral root on each first-order lateral root.

3.3. Nitrogen

As described in a previous study (Lu, et al., 2024b), nitrogen from soil can be taken up through two transporter systems: high (HATS, *µ*mol*/*(m^2^ · day), eq. S.8) and low (LATS, *µ*mol*/*(g root DW · day), eq. S.9) affinity systems. Besides, the previous experiment (Siddiqi et al., 1989) suggested that in addition to the HATS and LATS, there was an independent negative feedback also regulating N uptake. In our model, *E_N_* (dimensionless, eq. 10) represents a negative feedback of plant N concentration on root N uptake by the combined transport systems. Based on previous studies, this negative feedback was added to N uptake in the model (Barillot et al., 2016; Bertheloot et al., 2011). For each root segment, Michaelis-Menten kinetics were used to simulate potential N uptake through the HATS, and a linear relation with soil N to simulate potential N uptake through the LATS. The final N uptake is co-determined by available soil N in the soil cell and the plant demand of N (eq. S.11, S.12).

$HATS\left( t \right)=\frac{I_{max}\times\left( SoilN\left( t \right)-Nmin \right)}{Km+\left( SoilN\left( t \right)-Nmin \right)}$	(S.8)

*LATS*(*t*) = *K*_2_ × (*SoilN*(*t*) − *Nmin*)	(S.9)

$E_{N}=e^{-P\times Nc\left( t \right)}$	(S.10)

*potNup*(*t,x,N*) = *E_N_* × (*HATS*(*t*) × *A*(*x,N*) + *LATS*(*t*) × *rootDW*(*x,N*))	(S.11)

$Nup\left( t,x,N \right)=min(potNup\left( t,x,N \right), \left( soilN\left( t \right)-Nmin \right)*\frac{potNup(t,x,N)}{\sum_{N} \sum_{x} potNup(t,x,N)}$)	(S.12)

$Nuptake\left( t,N \right)=\sum_{x} Nup(t,x,N)$	(S.13)

*Nmin* (*µ*mol*/*L) is the minimum soil N concentration for nitrogen uptake. *I_max_* ( *µ*mol*/*(m^2^ · day)) is the maximum influx rate of N. *Km* (*µ*mol*/*m^3^) is the soil nitrogen concentration at half *I_max_*. Both parameter values were derived from York et al. (2016). *K*_2_ (*µ*mol*/*(g · day)) is the constant rate of *LATS* activity, which was derived from Pace and McClure (1986). *A* (m^2^) is the surface area of an individual root segment and *rootDW* (g) is the dry weight of the individual root segment. *P* (dimensionless) is the coefficient for the negative feedback of plant N concentration on root nitrogen uptake. This value was manually adjusted to avoid the maximum plant Nc exceeding 0.05g N/g DW. *Nc* (g N*/*g DW) is the whole plant nitrogen concentration. The x represents the order of the root segments, and the N represents the plant number.

After N uptake, two purposes of plant N are considered in the model. Firstly, the structural N (*structN*, g) for each organ (i.e. internode, grain and root) is considered (eq. S.14-S.16), which are the fixed fractions (*fN*, g N/g biomass) of organ biomass. This type of N cannot be remobilized.

$avaliableN\left( t \right)=avaliableN\left( t-1 \right)+Nuptake(t)$	(S.14)

*potstructN*(*i,t*) = fN*(i)* × growth(*i, t*)	(S.15)

$structN\left( i,t \right)=min(potstructN\left( i,t \right),avaliableN(t)\frac{potstructN(i,t)}{\sum_{i} potstructN(i,t)})$	(S.16)

where *i* represents the plant organs. The *availableN* (g per plant) represents the total amount of available N in the plant. The *potstructN* (g per plant organ) represents the demand for structural N per plant organ.

Besides the structural N in the plant organ, the remaining N is used for photosynthesis in the photosynthetic organs (i.e., leaf and internode). The organ demand for photosynthetic nitrogen (*DNareaphoto*, g/m2) is the nitrogen demand as a function of the light gradient in the canopy and the target nitrogen concentration (LN0, g/m2) of fully lit leaves (eq. S.17) (Hikosaka et al., 2016). This part of nitrogen can be redistributed.

*DNarea_photo_*(*i,t*) = *LN*0 × *Fabs^kNkL^*	(S.17)

$SPN(i,t)=min({DNarea}_{photo}\left( i,t \right), avaliableN*\frac{\frac{potphotoN\left( i,t \right)}{\sum_{i} potphotoN\left( i,t \right)}}{A_{i}})$	(S.18)

Where *A_i_* represents surface area of an individual photosynthetic organ (i.e. individual leaf or internode). *Fabs* is the fraction of incoming light absorbed by that organ. *kNkL* is the ratio between nitrogen and light extinction coefficients, and set equal to 0.368 (Hikosaka et al., 2016).

3.4. Tillering rule in the wheat model

A wheat plant produces multiple tillers during its growth, which all carry leaves. In this model, leaf number influences the shoot-derived auxin, which makes the shoot-derived auxin depend on tiller number. Based on the previous study, the outgrowth of a tiller happens when the number of phytomers is larger than a threshold value (J_ss_, Evers et al., 2010, eq. S.19). The threshold value is a function of the daily carbon source: sink ratio (eq. S.19). To further account for the variation of the tillers caused by shading, we introduce an estimated probability to calculate the possibility of a new tiller outgrowth (Wang et al., 2025).

$J_{ss}=\frac{1}{k+ss(t)}+dom$	(S.19)

$ss\left( t \right)=\frac{growth(t)}{\sum_{i} sinkstrength(i,t)}$	(S.20)

where k is the coefficient, and the dom represents the minimum phytomer between the apex and the top branch.

3.5. Soil

The soil is composed of voxels of size 0.1×0.1×0.1 m. Simulated soil depth is set to 2 m. Nitrogen is uniformly distributed at the beginning of the simulation. Periodic boundaries are applied, which means that root segments that grow beyond the later soil boundary enter the soil from the opposite site (de Vries et al., 2021).

3.6. Light

To calculate light absorption by the plant, parallel rays of light with specific azimuth and zenith angles were used to approximate the angular distribution of solar radiation in the sky, assuming clear sky conditions and computing total solar radiation as a function of day of year and latitude (Morales et al., 2025).

4. **Model evaluation scenario – fixed biomass allocation to the roots**

First, we tested how the newly implemented auxin-related mechanisms influence root architecture. The amount of biomass allocated to root growth each day was fixed to a constant ratio of root sink strength per day (0.7), to be able to analyze the effect of auxin on lateral root outgrowth and root elongation, without feedback effects on root biomass causing additional root architectural differences. The model was run under high nitrogen (HN, 3000 µmol/L) and low nitrogen (LN, 30 µmol/L) conditions. We first changed the input parameters one by one: [IAA]_Shoot_ (0.5, 1, 2 a.u. per day) and [IAA]_root_ (0.5, 1, 2 a.u. per day) in order to test how shoot or root-produced auxin influences root architecture. Then, we fixed the probabilities for lateral root initiation (P_init_) and emergence (P_emerge_), as well as the coefficient modifying potential root elongation rate (A_auxin_) to 1.0 to test how the newly implemented functions influence root architecture. The model was run for 20 days with 200 replications. For the root number at each soil layer, we chose the second axial root as it is less affected by seed nitrogen than the first emerged axial root. Also, at the early developmental stage, the rest of the axial root was still too short to provide fair observations. The outputs included root numbers on the second axial root and total root length per soil layer.

5. **Testing scenario**

When extremely low IAA is transported to cell division and cell elongation zones, the cell activity cannot be maintained (Wang et al., 2023). Therefore, the potential root elongation rate can be reduced. To account for this, a threshold (*V_auxin_*, a.u.) of *[IAA]_func_* is established, and beyond this threshold, we assume the potential root elongation rate is 0.

To test the effect of the V_auxin_ on plant biomass, we ran simulations with two different V_auxin_ values (1.5 and 1) under three low N conditions (10, 30 and 300 umol/L) for 70 days. In this scenario, we enabled the shoot part of the model and simulated light interception and N-dependent photosynthesis to drive biomass production. We ran the simulations 10 times to account for the randomness. After that, we recorded the relative total biomass (eq. 15).

6. **Supplementary results**
   6.1. Evaluation scenario- fixed biomass allocation to the roots

When increasing the strength of potential shoot auxin production rate, root number and root length increased across soil layers (Fig. S6A and B). Potential auxin production rate by roots had a smaller effect on root number and root length than when auxin was produced by shoots (Fig. S6C and S6D). High N increased root length in the top two soil layers compared with low N; however, it decreased root length in the deeper soil layers (Fig. S6A and S6C). Fewer lateral roots on the second emerged axial root were found in the top soil layer at high N, while more lateral roots formed in deeper soils compared with low N (Fig. S6B and S6D). The higher lateral root number under low N than under high N in the top soil layer was due to an effect on lateral root initiation, caused by higher total auxin at the early developmental stage (up until day 12) under low N than under high N (Fig. S6).

When fixing the potential root elongation rate coefficient (A_auxin_) to 1, root length and number were both reduced in all soil layers (Fig. S7A and S7D) with a slightly stronger decrease towards the deeper soil layers. With the probability for lateral root initiation fixed (P_init_ = 1), root number and length increased in the top two soil layers. There was no clear decrease or increase in the deeper soil layers (Fig. S7B and S7E). With a fixed probability for lateral root emergence (P_emerge_ = 1), root length and number increased across all the soil layers except in the deepest soil layer containing roots (Fig. S7C and S7F), also with a stronger increase towards the deeper soil layers.

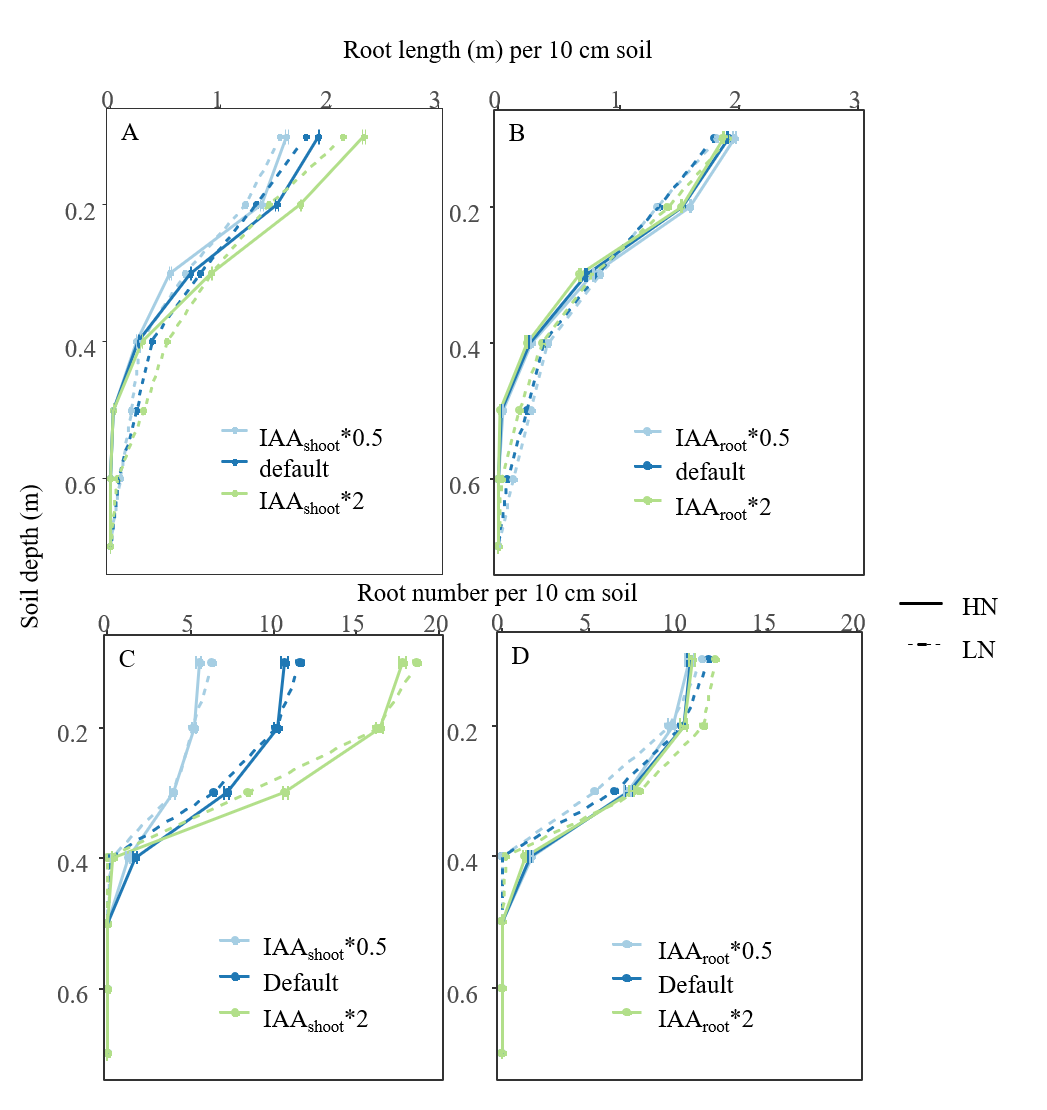

Fig. S7 Simulated root length and lateral root number on the second axial root across the soil profile. The model was run under high N (3000 μmol/L) and low N (30 μmol/L) for 20 days with different values of the potential shoot or root auxin production rate parameter ([IAA]_shoot_ or [IAA]_root_ times 0.5, 1.0, and 2.0). The values represent mean ± se (n=200).

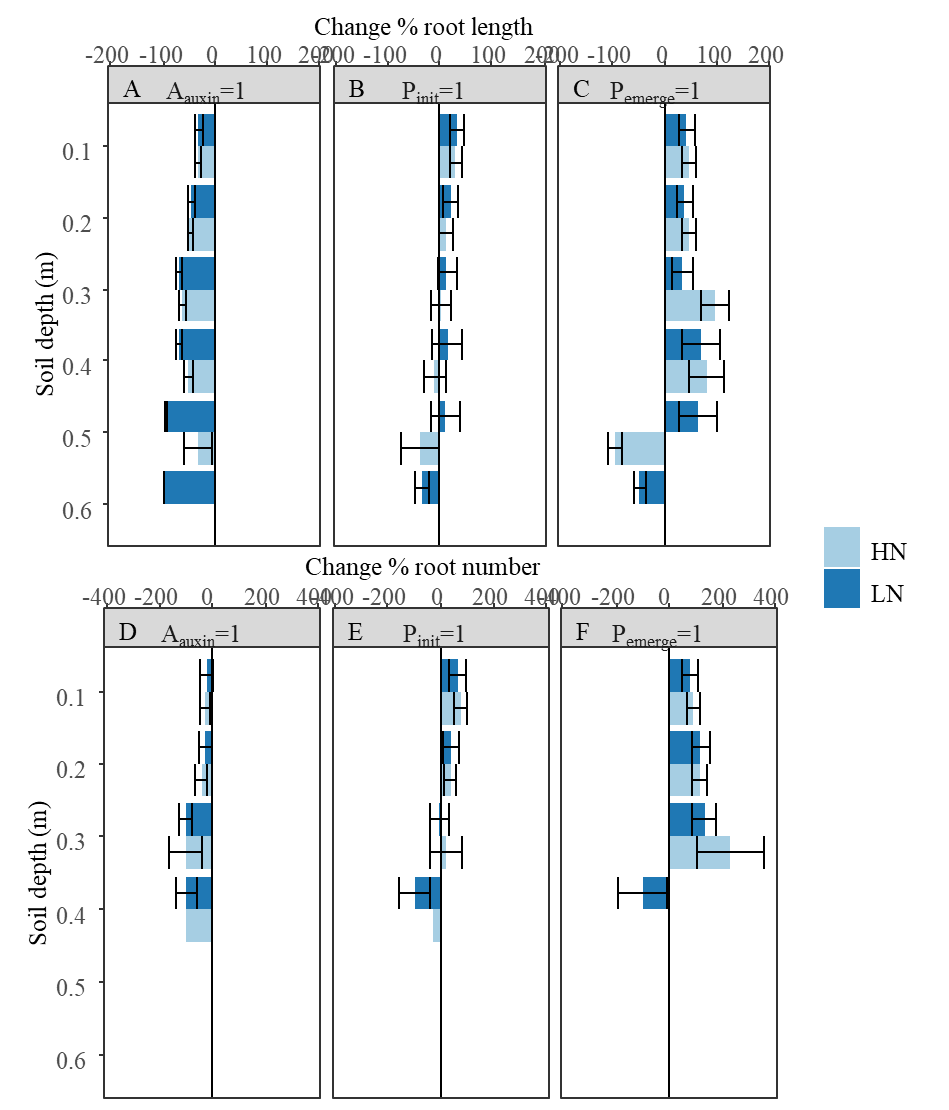

Fig. S8 Simulated changes in root length and lateral root number across the soil profile. Since there was no root beyond 60 cm soil depth at this growth stage, the soil profile from 0 to 60 cm was showed here. The model was run under high N (3000 μmol/L) and low N (30 μmol/L) for 20 days with disabling different auxin related functions (A_auxin_, P_init_, P_emerge_) in the model. A_auxin_ represents the modifier of the potential root elongation rate, which a function of total auxin in the root meristem. P_init_ represents the probability of lateral root initiation, which is a function of total auxin. P_emerge_ represents the probability of lateral root emergence, which is the function of shoot-derived auxin. The values represent mean ± se (n=200).

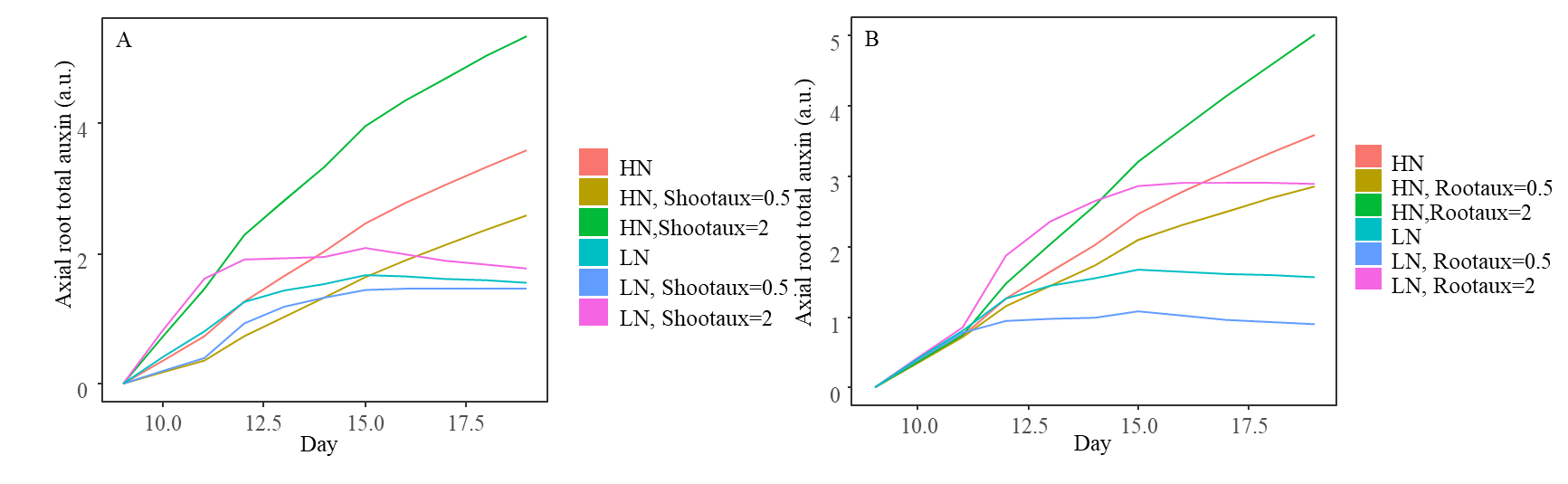

Figure. S9 Effective auxin in axial root with shoot/root auxin levels under high N (3000 umol/L) and low N (30umol/L) conditions.

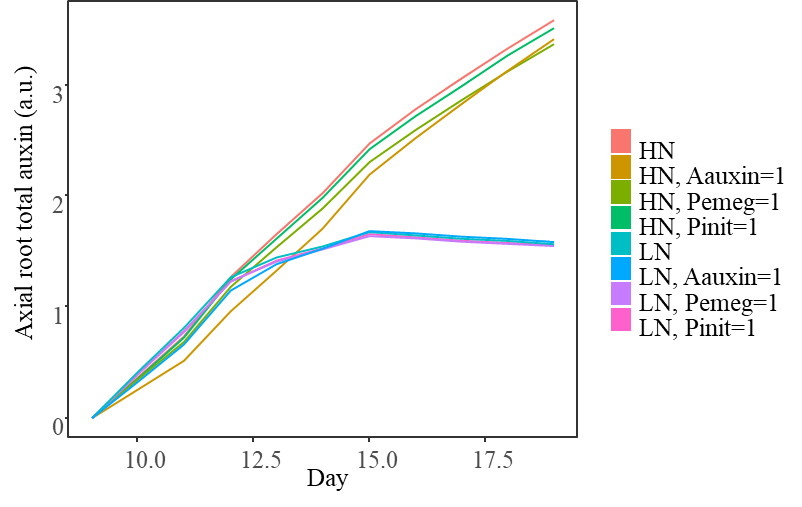

Figure S10 Effective auxin in the axial root of disabled auxin-related processes under high N (3000 umol/L) and low N (30 umol/L).

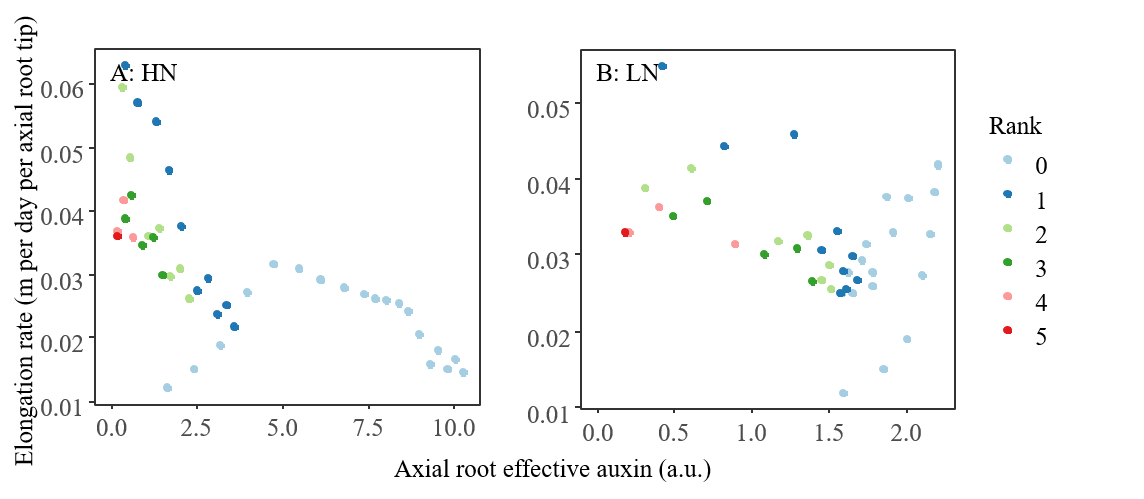

Fig. S11 Simulated actual axial root elongation rate vs axial root effective auxin under high N (HN, 3000 umol/L) and low N (LN, 30 umol/L). Rank 0 represents the first emerged axial root (primary root). Rank 5 represents the latest-emerged axial root at day 20.

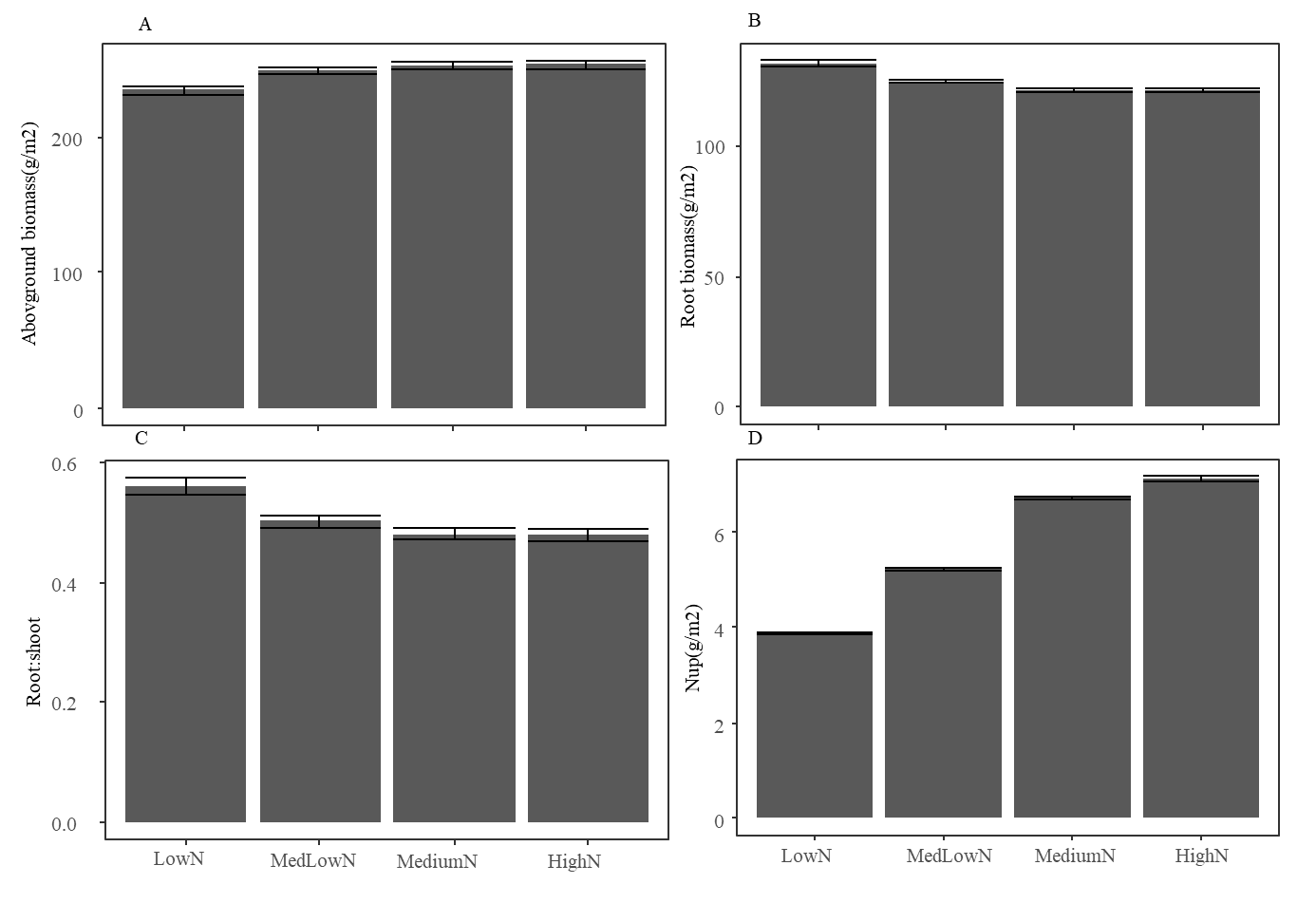

Fig. S12 Simulated plant characteristics (absolute values) across multi-N levels at planting density of 343 plants/m^2^. The LowN, MedLowN, MediumN and HighN represent total N levels in simulation: 500, 1000, 2000, and 3000 µmol/L. The simulations begin at the 77^th^ day of the year and run for 49 days after sowing. Values are mean ± SE (n=10).

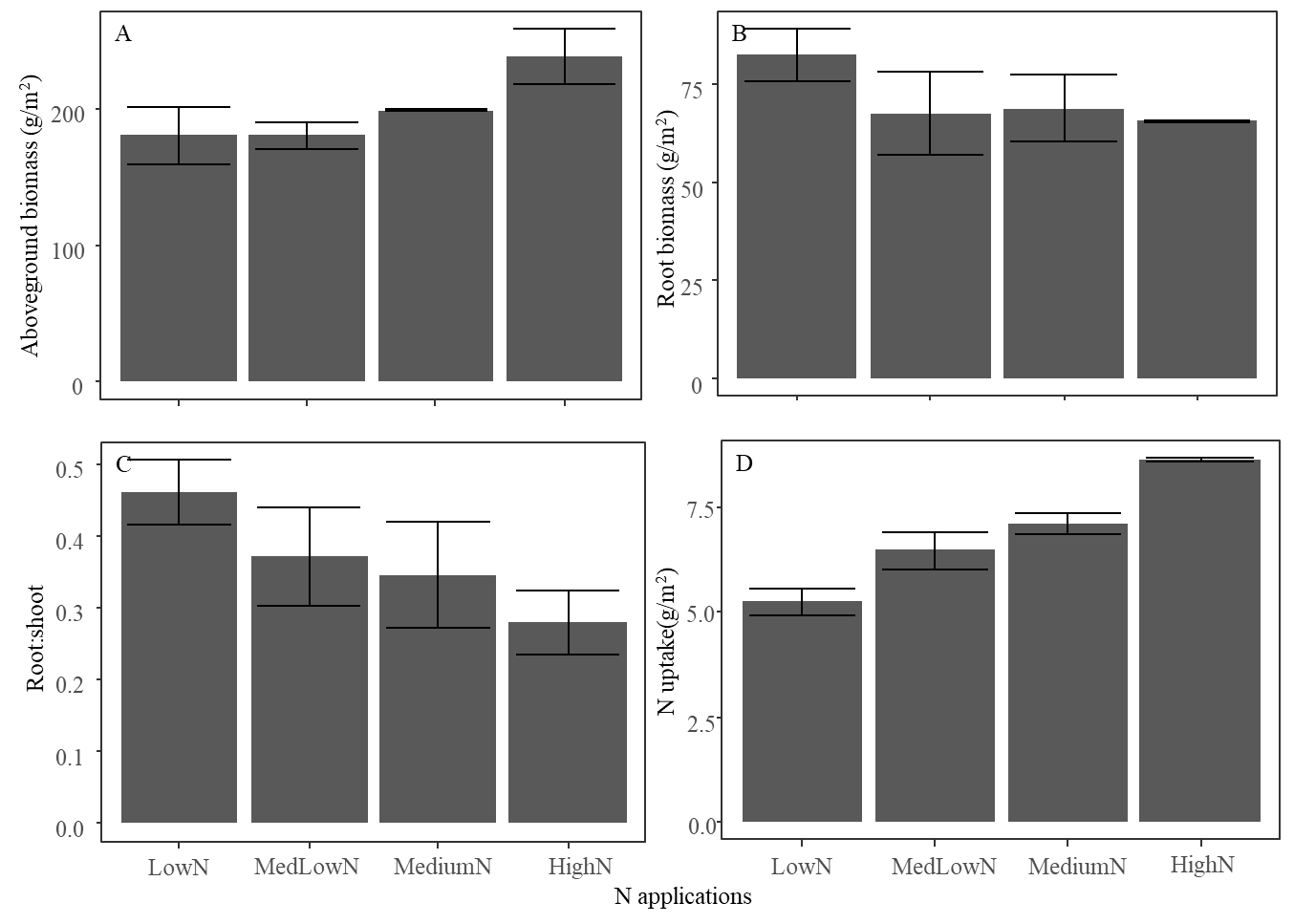

Fig. S13 Observed plant characteristics (absolute values) across multi-N levels at planting density of 343 plants/m^2^. The LowN, MedLowN, MediumN and HighN represent applied N in the soil: 0, 2, 4, and 6 g/m2. Values are mean ± SE (n=3).

6.2. Exploring the effect of local/systemic responses

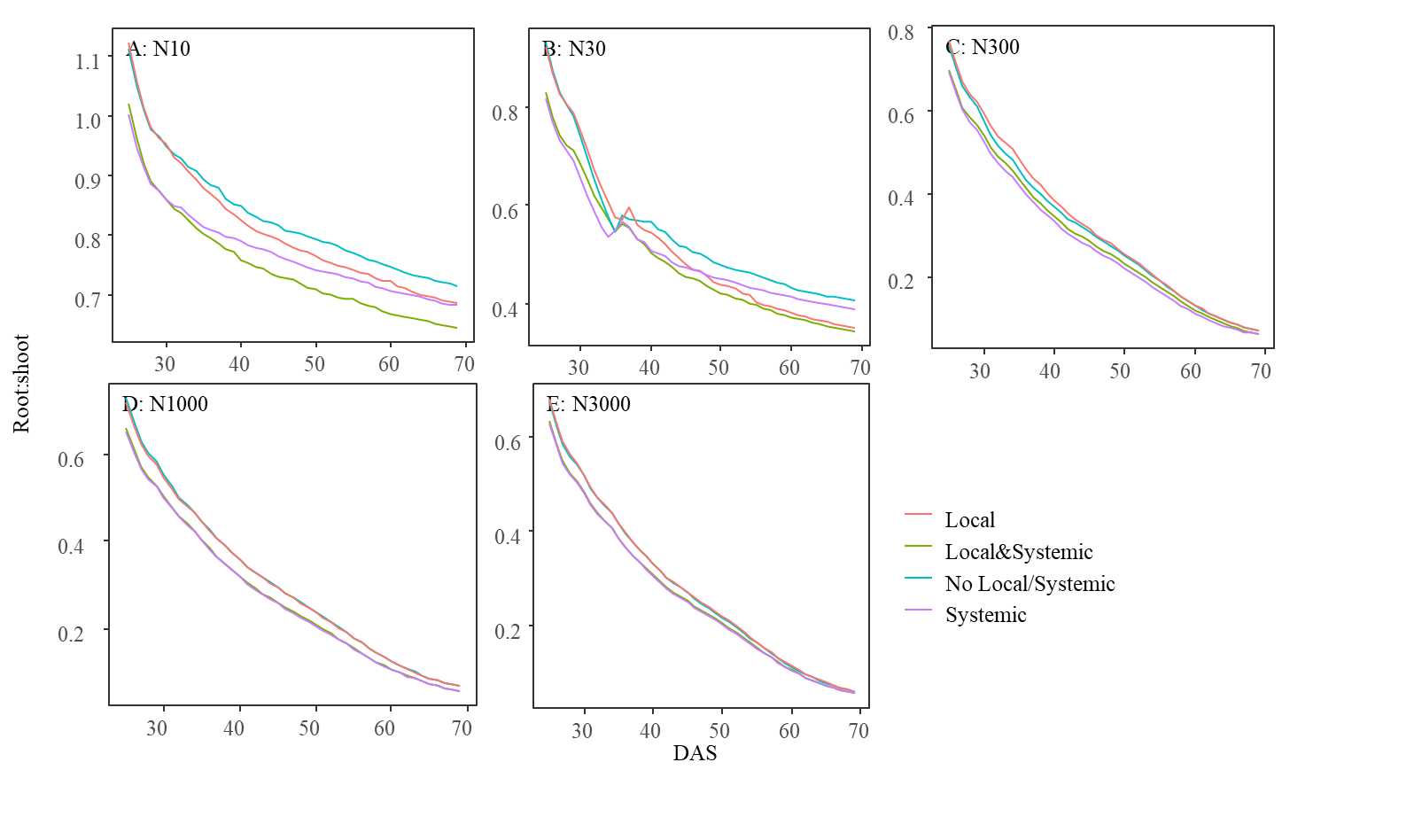

Figure S14 Simulated root-to-shoot ratio over days across N levels. In the simulations, whole plant feedback and photosynthesis were enabled.

6.3. Testing scenario

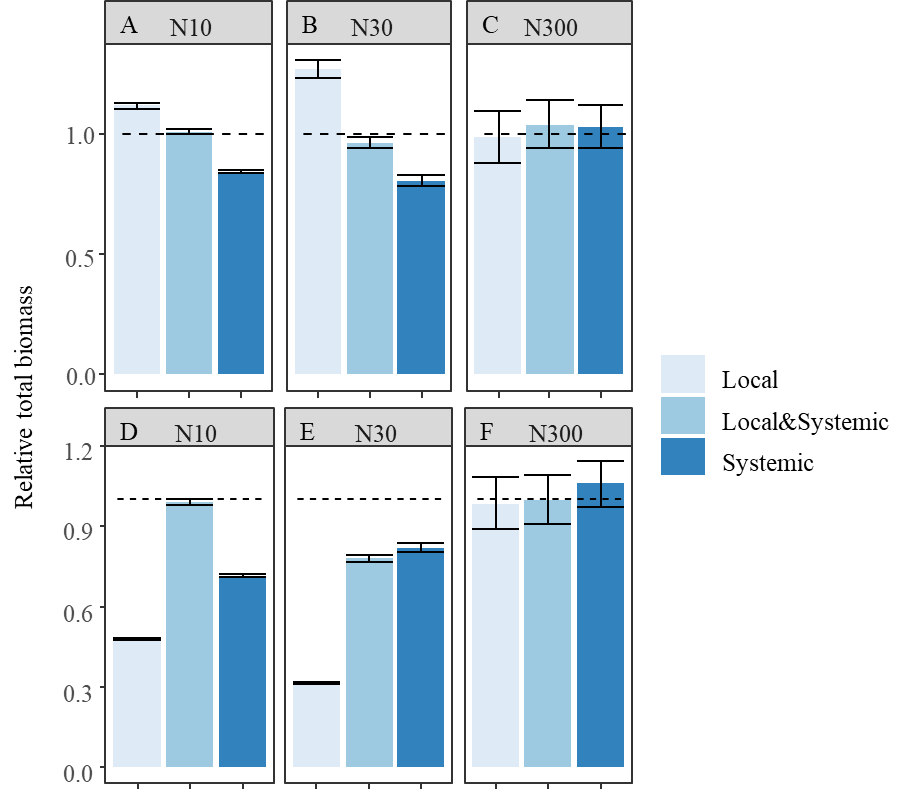

Fig. S15 Simulated total biomass for different response types, relative to total biomass without either response. The Panels A, B, C represent the simulations with *V_auxin_* equal to 1. The Panels D, E, F represent the simulations with *V_auxin_* equal to 1.5. The simulations were run for 70 days. The dotted line represents a relative N uptake of 1. ‘Local’, ‘Local & Systemic’, and ‘Systemic’ indicate which responses to N were enabled in simulation. N10, N30 and N300 represent the initial soil N at 10, 30, and 300 µmol/L. The values represent mean ± SEs (n=10).

Supplementary tables

| Table S1: Definitions of auxin-related variables | | |  |
| --- | --- | --- | --- |
| Equation# | Variables | Definition | Unit |
| 3 | [IAA]_func_ | The auxin can move to cell elongation and division zones in the root tip and contribute to changes in root elongation and lateral root initiation | a.u. |
| 4 | P_emerge_ | The probability of a lateral root emergence |  |
| 5 | A_auxin_ | A modifier of the potential root elongation rate as function of total auxin in the root meristem |  |
| 6 | EL_auxin_ | Potential root elongation rate driven by total auxin in the root meristem |  |
| 7 | P_init_ | The probability of lateral root initiation |  |
| 8 | γ | The relative auxin concentration at the root tip caused by translocation in response to plant internal N |  |
| 9 | [IAA]_shoot_ | The rate of shoot produced auxin transported to the roots at given plant N concentration | a.u. |
| 10 | F_rp_ | The multiplier of potential root-produced auxin to calculate the actual auxin produced in the root meristem in response to the external nitrate conditions |  |
| 11 | [IAA]_root_ | The rate of root produced auxin at given external N conditions | a.u. |
| 12 | F_reduction_ | The multiplier of the combined rate for IAA reduction to calculate the rate of auxin reduction in the root meristem in response to the external nitrate conditions |  |
| 13 | δ_exN_ | The rate of auxin transport out of the meristem | per day |

| Table S2: Auxin-related parameter values | | | |  |  |
| --- | --- | --- | --- | --- | --- |
| # | Equation | Parameters | Definition | Values | Reference |
| 4 | P_emerge_=1-a_S_ exp(-b_S_ [IAA]_shoot_ ) | a_S_ | the magnitude factor of the exponential curve | 1,033 | Reed et al., 1998 |
|  |  | b_S_ | steepness of the exponential curve | 0,212 | Reed et al., 1998 |
| 5 | A_auxin_=a_T_exp(b_T_[IAA]_eff_)+1 | a_T_ | the magnitude factor of the exponential curve | 1,75 | Reed et al., 1998 |
|  |  | b_T_ | steepness of the exponential curve | -0,343 | Reed et al., 1998 |
| 6 | EL_auxin_= EL*A_auxin_ | EL | potential elongation rate (m/(m day)) | 23 | Sharma & Ghildyal, 1987 |
| 7 | P_init_=a_i_[IAA]_eff_exp(b_i_[IAA]_eff_)+c_i_ | ai | Fitted coefficients | 0,161 | Ivanchenko et al., 2010 |
|  |  | bi | Fitted coefficients | -0,128 | Ivanchenko et al., 2010 |
|  |  | ci | baseline probability of the lateral root initiation when auxin is high | 0,491 | Ivanchenko et al., 2010 |
| 8 | γ=a_inN_-b_inN_ Nc_min_, Nc_total_<Nc_min_ | ainN | relative auxin concentration at the root tip translocated from shoot when plant N concentration is 0 | 1,58 | Tian et al., 2008 |
|  | γ=a_inN_-b_inN_ Nc_total_, Nc_min_< Nc_total_<Nc_max_ | binN | the decreased rate of relative auxin concentration at the root tip translocated from shoot with increasing plant N concentration | 26,68 | Tian et al., 2008 |
|  | γ=a_inN_-b_inN_ Nc_max_, Nc_total_>Nc_max_ | Ncmin | minimum plant N concentration (g/DW g) | 0,0216 | Tian et al., 2008 |
|  |  | Ncmax | maximum plant N concentration (g/DW g) | 0,0478 | Tian et al., 2008 |
| 9 | [IAA]_shoot_=γ[IAA]_s,pot_ | [IAA]_s.pot_ | potential rate of shoot produced auxin translocated to the root meristem (a.u./day) | 1 |  |
| 10 | F_rp_=Max(0.18,1-b_rp_ Nexternal) | b_rp_ | the coefficients of the relative root-produced IAA in response to external N concentration | 0,00082 | Liu et al.,2010 |
| 11 | [IAA]_root_=F_rp_[IAA]_r.pot_ | [IAA]_r,pot_ | potential auxin produced in each axial root meristem (a.u./day) | 1 |  |
| 12 | F_reduction_=Max(0,1-b_reduction_ Nexternal) | b_reduction_ | the coefficient of the relative IAA reduction rate in response to external N concentration | 0,001 | Liu et al.,2010 |
| 13 | δ_exN_=F_reduction_δ | δ | IAA reduction rate in the root meristem | 0,43 | Grieneisen et al., 2007 |

| Table S3: Plant level parameter values | |  |  |
| --- | --- | --- | --- |
| **Parameter** | **Description** | **Example value** | **References** |
| **Shoot parameters** | | | |
| **shoot architecture/development related parameters** | | | |
| LeafNum | Final number of vegetative phytomers | 10 | Tilley et al., 2019 |
| phyllotaxis | Angle between consecutive leaves along a stem for the lower phytomers | 137 | Default values from original model |
| maxIntwidth | Maximum internode width (m) | 0,005 | Default values from original model |
| specificIntLength | Internode length per mg of biomass (m/g) | 0,6 | Indirect estimation |
| Lowrank | Number of Internode without enlongation | 4 | Tilley et al., 2019 |
| maxWidth | Location on blade of maximum width (fraction distance from tip) | 0,7249 | Default values from original model |
| shapeCoeff | Leaf shape coefficient (0 = rectangular, 1 = pinched) | 0,2027 | Default values from the original model |
| leafAngle | Angle between leaf blade base and stem (90 = horizontal) | 40 | Default values from the original model |
| leafCurve | Angle between tangent at blade base and tangent at blade tip (0 = flat blade, no curvature) | 46 | Default values from the original model |
| lwRatio | Leaf blade length / width ratio | 27 | Default values from the original model |
| specificsheathlength | Sheath length per mg of biomass (m/g) | 2.5 | Default values from the original model |
| Maxtiller | maximum tillers | 6 | Wang et al., 2025 |
| plastochron | Thermal time interval between successive leaf initiation at the apex (°Cd) | 43 | Default values from the original model |
| phyllochron | Thermal time interval between appearance of two leaves (^o^Cd) | 86 | Default values from the original model |
| tb | Base temperature for plant development (°C) | 0 | Default values from the original model |
| **C sink related parameters** | | | |
| wmax | Potential biomass of grain (g) | 3 | Default values from the original model |
| wmaxInt | Potential biomass of an internode (g) | 0.266 | Default values from the original model |
| wmaxLeaf | Potential biomass of a leaf (g) | 0.266 | Default values from the original model |
| wmaxRoot | Potential biomass of root system (g) | 1 | Estimated from Wang et al., 2024 |
| te | Growth duration of grain (°Cd) | 800 | Default values from the original model |
| teInt | Growth duration of an internode (°Cd) | 182 | Default values from the original model |
| teRoot | Growth duration of root system (°Cd) | 800 | Indirect estimation |
| teLeaf | Growth duration of a leaf (°Cd) | 220 | Default values from the original model |
| sheathFraction | Fraction of leaf biomass partitioned to the sheath | 0,05 | Default values from the original model |
| **C source related parameter** | | | |
| seedMass | Endosperm mass of the seed the plant grows from initially (g) | 0.025 | Default values from the original model |
| α | Initial light use efficiency (initial slope of light response curve) | 0,015 | Zheng et al., 2021 |
| λ | Maximum photosynthesis under non-limiting leaf nitrogen (umol /(m^2^ s)) | 23.6 | Zheng et al.,2021 |
| **N related parameters** | | | |
| LN0 | Target nitrogen content of leaves at top of canopy (g/m2 of leaf) | 3 | Indirect estimation |
| kNkL | The ratio between nitrogen and light extinction coefficients | 0.368 | Hikosaka et al., 2016 |
| fNgrain | Grain N concentration (g N/ DW g) | 0.03 | Haberle et al., 2008 |
| fNstem | Stem structural N concentration(g N/ DW g) | 0.01 | Personal discussion with wheat modeler |
| fNroot | Root N concentration (g N/ DW g) | 0.005 | Personal discussion with wheat modeler |
| **Root Parameter** | | | |
| **Root biomass related parameters** | | | |
| a_r_ | Coefficient for root to leaf partitioning fraction normalized by the minimum root-to-leaf ratio in response to plant N concentration | 2.747 | Lu et al., 2024 |
| b_r_ | Coefficient for root to leaf partitioning fraction normalized by the minimum root-to-leaf ratio in response to plant N concentration | 100 | Lu et al., 2024 |
| **Root architecture related parameters** | | | |
| initD | Initial root Diameter (m) | 0.001 | Dimattia et al. 2025 |
| RTD | Root tissue density (g/cm3) | 60 | Nakhforoosh et al., 2014 |
| RootNum | Maximum number of seminal roots | 30 | Wang et al., 2025 |
| ER | Rate of emergence of seminal roots (per °Cd) | 0,056 | Page et al., 2014 |
| rootseg | Root segment length (m) | 0,003 | Page et al., 2014 |
| Dfine | Fine root density (m fine roots/m first order lateral root) | 8,12 | Estimation from Chen et al., 2014 |
| RDM | Ratio between diameter of mother and daughter root | 0.12 | Page et al., 2014 |
| angleAVG | Insertion angle of lateral roots (^o^) | 60 | Default values from the original model |
| angleVAR | Standard deviation of angleAVG | 40 | Default values from the original model |
| InitAngle | Initial axial root angle with vertical stem (^o^) | 60 | Oyanagi 1994 |
| InitAngleVAR | Standard deviation of InitAngle | 20 | Indirect estimation |
| MCP | Random root deviation from it original orientation based on mechanical constraints (radial degree/m) | 5 | Default values from original model |
| Groot | Base rate of gravitropism | 0,05 | Default values from original model |
| **Root physiological related parameter** | | | |
| Imax | Maximum nitrogen influx rate under saturate soil condition for high affinity nitrogen transporter (umol/m2/day) | 25000 | York et al., 2016 |
| Km | External nitrogen at half of Imax (umol/L) | 50 | York et al., 2016 |
| K_2_ | Linear parameter for low affinity nitrogen transporter(umol/g/day) | 1,46 | Pace and McClure, 1986 |
| P | The coefficient for the negative feedback of plant N concentration on root nitrogen uptake | -80,00 | Indirect estimation |
| Note: since this model is a further development from an existing model (see Lu et al., 2024, De Vries et al. 2021, Evers, et al., 2010) and this existing model includes multiple species, some parameter names are different in the code. We therefore provide notations of the name used in this manuscript behind the actual name of the parameters in the code | | | |

| Table S4 Environment parameter values | | |  |  |
| --- | --- | --- | --- | --- |
|  | Evaluation scenario 1 | Evaluation scenario 2 | Research scenario 1 | Research scenario 2 |
| Location information |  |  |  |  |
| altitude | NA | 52 | NA | 52 |
| Temperature | 10.7+7.55*sin(2*pi*(dayofyear-111)/365) | 10.7+7.55*sin(2*pi*(dayofyear-111)/365) | 10.7+7.55*sin(2*pi*(dayofyear-111)/365) | 10.7+7.55*sin(2*pi*(dayofyear-111)/365) |
| Soil N concentration | Multi-levels | Multi-levels | Multi-levels | Multi-levels |
| Managements | | | | |
| row_distance | 0,2 | 0,12 | 0,2 | 0,2 |
| plant_distance | 0,2 | 0,025 | 0,2 | 0,2 |
| sowing date | 90 | 77 | 90 | 90 |
